# Cell-type-specific whole-genome landscape of ΔFOSB binding in nucleus accumbens after chronic cocaine exposure

**DOI:** 10.1101/2022.08.27.505547

**Authors:** Szu-Ying Yeh, Molly Estill, Casey K. Lardner, Caleb J. Browne, Angelica Minier-Toribio, Rita Futamura, Katherine Beach, Catherine A. McManus, Song-jun Xu, Shuo Zhang, Elizabeth A. Heller, Li Shen, Eric J. Nestler

## Abstract

The ability of neurons to respond to external stimuli involves adaptations of gene expression. The transcription factor, ΔFOSB, is important for the development of drug addiction, however, its gene targets have not been identified. Here we use CUT&RUN to map the genome-wide enrichment of ΔFOSB binding in the two major neuronal cell types of the nucleus accumbens, a key brain reward region, after cocaine exposure. The binding landscape shows that the majority of ΔFOSB peaks occur outside of promoter regions, including intergenic regions, and are surrounded by epigenetic marks indicative of active enhancers. BRG1, the core subunit of the SWI/SNF chromatin remodeling complex, overlaps with ΔFOSB peaks, consistent with earlier studies of ΔFOSB’s interacting proteins. In addition, *in silico* analyses predict that ΔFOSB cooperatively regulates gene expression with homeobox and T-box transcription factors. These novel findings uncover key elements of ΔFOSB’s molecular mechanisms in transcriptional regulation.

## Introduction

The powerful functions of the brain, including cognition, emotional regulation, learning, and memory, depend in part on the ability of neurons to alter their gene expression, and downstream activity and connectivity, upon encountering environmental challenges. With this power comes vulnerability, however: maladaptations can occur in response to detrimental stimuli, such as drugs of abuse, leading to syndromes of addiction in vulnerable individuals.

Modification of gene expression involves crosstalk among chromatin remodeling complexes, histone-modifying enzymes, and transcription factors (TFs). TFs are proteins that bind to specific nucleotide sequences, called response elements, across the genome. One such transcription factor, ΔFOSB, has been implicated as one important mechanism through which repeated exposure to a drug of abuse induces addiction-related behavioral abnormalities. ΔFOSB accumulates in neurons of the nucleus accumbens (NAc), a central node of the brain’s reward circuitry, in response to repeated exposure to virtually all classes of drugs of abuse in rodents, with such accumulation also documented in postmortem samples of humans with substance use disorders (SUDs)^1-3^. Further, such induction of ΔFOSB is known to display cell-type specificity. Approximately 95% of all NAc neurons are medium spiny projection neurons (MSNs) that are classified into two main subpopulations based on their predominant expression of either D1 or D2 dopamine receptors. These two subpopulations in NAc encode distinct—in some cases opposite—types of information during a course of drug administration^4,5^. All drugs of abuse induce ΔFOSB selectively in D1 MSNs, whereas opioids uniquely induce ΔFOSB equally in both D1 and D2 MSNs^6^. Overexpressing ΔFOSB selectively in D1 MSNs, by use of viral-mediated gene transfer or of inducible genetic mutant mice, increases an animal’s sensitivity to the rewarding effects of cocaine and certain other drugs of abuse, while antagonizing ΔFOSB action via expression of a dominant-negative antagonist exerts the opposite effect^1,2^. The consequences of ΔFOSB induction in D2 MSNs have not yet been characterized. Moreover, ΔFOSB also exerts different effects on the neurophysiological properties of D1 vs. D2 MSNs^7^.

Several prior studies have sought to identify target genes for ΔFOSB in NAc MSNs by examining expression of putative targets on a candidate gene basis or through more open-ended analysis, most recently by RNA-sequencing (RNA-seq)^8,9^. However, it has been far more difficult to interrogate direct genomic targets of ΔFOSB due to technical limitations. An earlier study used a ChIP-chip approach: chromatin immunoprecipitation of ΔFOSB followed by a promoter array which by definition is severely limited due to its focus on promoter regions only^10^. This study also analyzed bulk NAc tissue, thus not providing cell-type-specific information. Moreover, it has not been technically possible to use ChIP followed by sequencing (ChIP-seq) to analyze ΔFOSB binding sites in the genome. This difficulty is indicative of challenges in mapping transcription factor binding in the brain in general, especially in a micro-dissected brain region such as the NAc.

To overcome these obstacles and explore the direct genomic targets of ΔFOSB, we utilized a cutting-edge adaptation of ChIP-seq named CUT&RUN (cleavage under targets and release using nuclease)^11^. By use of this approach, we succeeded in obtaining high quality maps of ΔFOSB binding across the genome selectively in D1 MSNs and in D2 MSNs under control conditions and in response to chronic cocaine exposure in both male and female mice. Our findings provide a powerful template for characterizing transcription factor binding in sorted neuronal populations and offer new insight into the cell-type- and sex-specific molecular actions of ΔFOSB in the NAc.

## Results

### The majority of ΔFOSB binding sites in the NAc occur outside promoter regions

To establish the new ΔFOSB CUT&RUN-seq methodology, we first utilized bulk NAc tissue punches from wild-type male mice dissected 24 hr after the last of 7 daily intraperitoneal (IP) injections of either saline or cocaine. Several thousand peaks of putative ΔFOSB binding were found in both groups (Fig. 1A & S1A), and >93% of the called ΔFOSB peaks contained AP1 sites – the response element to which all FOS family proteins, complexed with their JUN family protein partners, canonically bind in other systems^12^. By contrast, only ∼20% of randomly-selected genomic regions contained AP1 sites (∼93% vs. ∼20% for random genomic regions). This confirms the ability of our approach to accurately capture bona fide ΔFOSB binding sites with great specificity. Chronic cocaine exposure induced a several-fold increase in the number of ΔFOSB peaks, with 5,352 peaks unique to cocaine condition vs. only 1,205 peaks unique to saline condition. An additional 7,315 peaks were shared between both conditions (Fig. 1B,C). These findings indicate that cocaine predominantly induces ΔFOSB binding to genomic targets in the NAc, consistent with cocaine induction of ΔFOSB expression in this brain region, whereas a smaller number of targets become depleted of ΔFOSB despite higher total nuclear levels of the protein.

**Figure 1.**
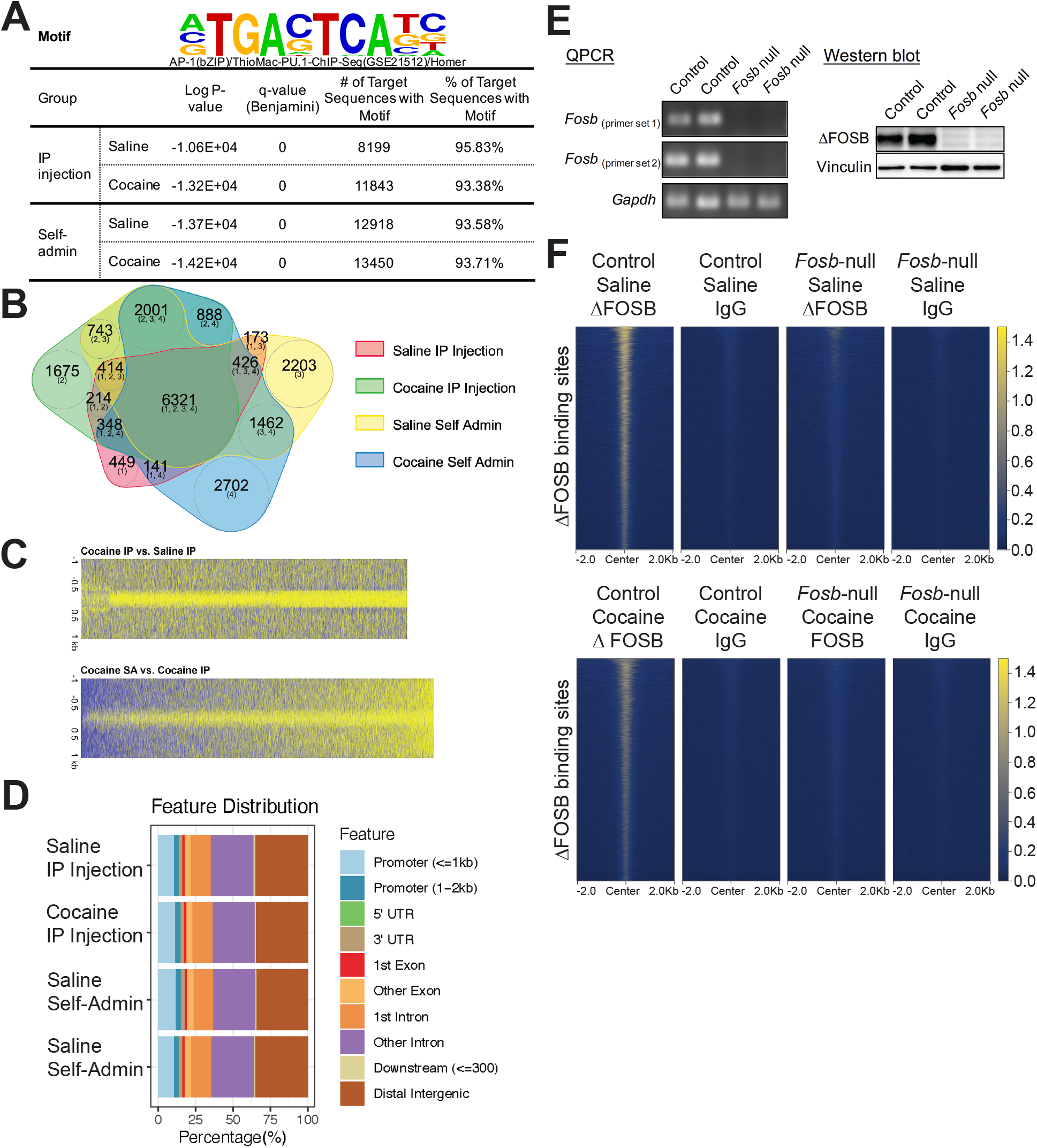
Establishment of CUT&RUN-seq for ΔFOSB in NAc. (A) Motif analysis of called ΔFOSB peaks in NAc of male mice. (B) Venn diagram analysis of called ΔFOSB peaks after repeated (7 d) cicine IP injections or chronic (10 d) cocaine self-administration and their saline controls. (C) Upper heatmap shows genomic regions where ΔFOSB binding is induced by repeated IP injections of cocaine vs. saline, and lower heatmap compares the effect of cocaine self-administration (SA) vs. IP injections. (D) Genomic distribution of ΔFOSB peaks in the NAc. (E) RNA and protein analysis of dorsal striatum from *Fosb-*null male mice. (F) Heatmap of ΔFOSB coverage at called ΔFOSB binding sides between control and *Fosb-*null groups.

We next identified genomic regions to which ΔFOSB binds under saline control and cocaine-treated conditions by annotating the peaks based on their location in the mouse reference genome (mm10). Surprisingly, only ∼15% of ΔFOSB peaks were found within putative promoter regions, i.e., −5,000 bp and +2,000 bp of known transcription start sites (TSSs). Roughly 1/3 of the peaks were located in intergenic regions and roughly half of the peaks were found within gene bodies (Fig. 1D). This distribution of ΔFOSB binding sites was equivalent under saline and cocaine conditions. While the preponderance of ΔFOSB binding sites outside promoter regions may seem surprising, a similar pattern has been observed for some transcription factors in cultured cells or peripheral tissues (see Discussion).

Rodents that volitionally self-administer cocaine are considered a more translational model of humans with substance use disorders, although we have shown that investigator- and self-administrated cocaine induce comparable levels of ΔFOSB in the NAc in rats and mice^6,13,14^. To determine whether ΔFOSB displays a similar binding landscape after cocaine self-administration, we performed ΔFOSB CUT&RUN-seq on bulk NAc dissections from wild-type male mice 24 hr after 10 d of intravenous cocaine or saline self-administration (Fig. S1C). The vast majority of identified peaks once again contained an AP1 motif (∼93% vs. ∼20% for random genomic regions) (Fig. 1A and S1B), and we found substantial overlap in ΔFOSB peaks in the self-administration vs. IP injection groups, although ∼2,500 peaks were unique to the self-administration datasets (Fig. 1B). The genomic distribution of ΔFOSB binding sites was also similar, with only a small portion (<15%) of the peaks present at promoter regions (Fig. 1D). Interestingly, despite the induction of comparable total levels of ΔFOSB protein in NAc in response to IP vs. self-administered cocaine, in general ΔFOSB binding was stronger under the self-administration conditions (Fig. 1C).

To confirm that the peaks identified from the ΔFOSB CUT&RUN-seq experiments are indeed bona fide ΔFOSB binding sites, we applied the same methodology to tissues from *Fosb*-null mice. To generate *Fosb-*null mice, we crossed *Sox2-Cre* mice to *Fosb-flox/flox* mice^13^. *Sox2-Cre;Fosb-flox/flox* (*Fosb-*null) mice and littermate controls (*Fosb-flox/+*) were given 7 daily cocaine or saline IP injections and analyzed 24 hr after the last injection, as before. To verify that the *Fosb-*null mice did not express ΔFOSB, we extracted RNA and protein from the dorsal striatum for qPCR and Western blotting, respectively. Δ*Fosb* transcript and ΔFOSB protein were not detectable in *Fosb-*null samples (Fig. 1E). We next performed ΔFOSB CUT&RUN-seq on NAc dissections from *Fosb-*null and control mice and found that only control samples displayed ΔFOSB binding sites (Fig. 1F), similar to those observed in wild-type mice, with a virtually complete loss of significant ΔFOSB peaks in NAc of *Fosb*-null mice. Additionally, CUT&RUN-seq utilizing non-immune IgG showed a similar loss of ΔFOSB binding sites. These crucial control data demonstrate that the ΔFOSB antibody used for CUT&RUN-seq captures ΔFOSB specifically and not other members of the FOS family or non-specific, artifactual binding. Although the *Fosb* gene expresses full-length FOSB in addition to its truncated splice variant ΔFOSB, both of which are recognized by the antibody used, ΔFOSB alone is detectable in NAc 24 hr after chronic cocaine or saline treatment^1^.

### ΔFOSB binding maps in D1 MSNs and D2 MSNs of the NAc

Given the very different patterns of ΔFOSB induction in D1 vs. D2 NAc MSNs in response to cocaine and other stimuli^6^, and the very different roles of these two neuronal subpopulations in cocaine action (see Introduction), we next set out to obtain genome-wide maps of ΔFOSB binding in these two MSN subtypes. Given the very different transcriptional effects of cocaine seen in NAc between the sexes^19^ and the very different gene expression changes seen in D1 and D2 MSNs upon ΔFOSB induction in the NAc of male vs. female mice^9^, we generated such ΔFOSB maps from NAc D1 and D2 MSNs of male and of female mice. To enable the selective isolation of nuclei from D1 and from D2 MSNs, we generated animals with GFP-labeled nuclei by crossing *D1-Cre* or *D2-Cre* mice with *Rosa-Sun1/sfGFP* mice. Nuclei from NAc D1 MSNs or from D2 MSNs were isolated by INTACT (isolation of nuclei tagged in specific cell types) approach and then subjected to ΔFOSB CUT&RUN-seq.

Comparable to bulk NAc experiments, thousands of ΔFOSB peaks were identified from NAc D1 and D2 MSNs, with >95% of the loci containing consensus AP1 motifs (∼95% vs. ∼20% for random genomic regions) (Fig. 2A & S2A-B). A majority (∼75%) of ΔFOSB peaks were shared between D1 and D2 MSNs in both males and females (Fig. 2B). The distribution of the peaks was equivalent to that seen in bulk NAc: only ∼15% of ΔFOSB binding sites were located at promoter regions, and the rest were located within intergenic regions and gene bodies (Fig. 2C). These features were equivalent between neurons isolated from male vs. female mice.

**Figure 2.**
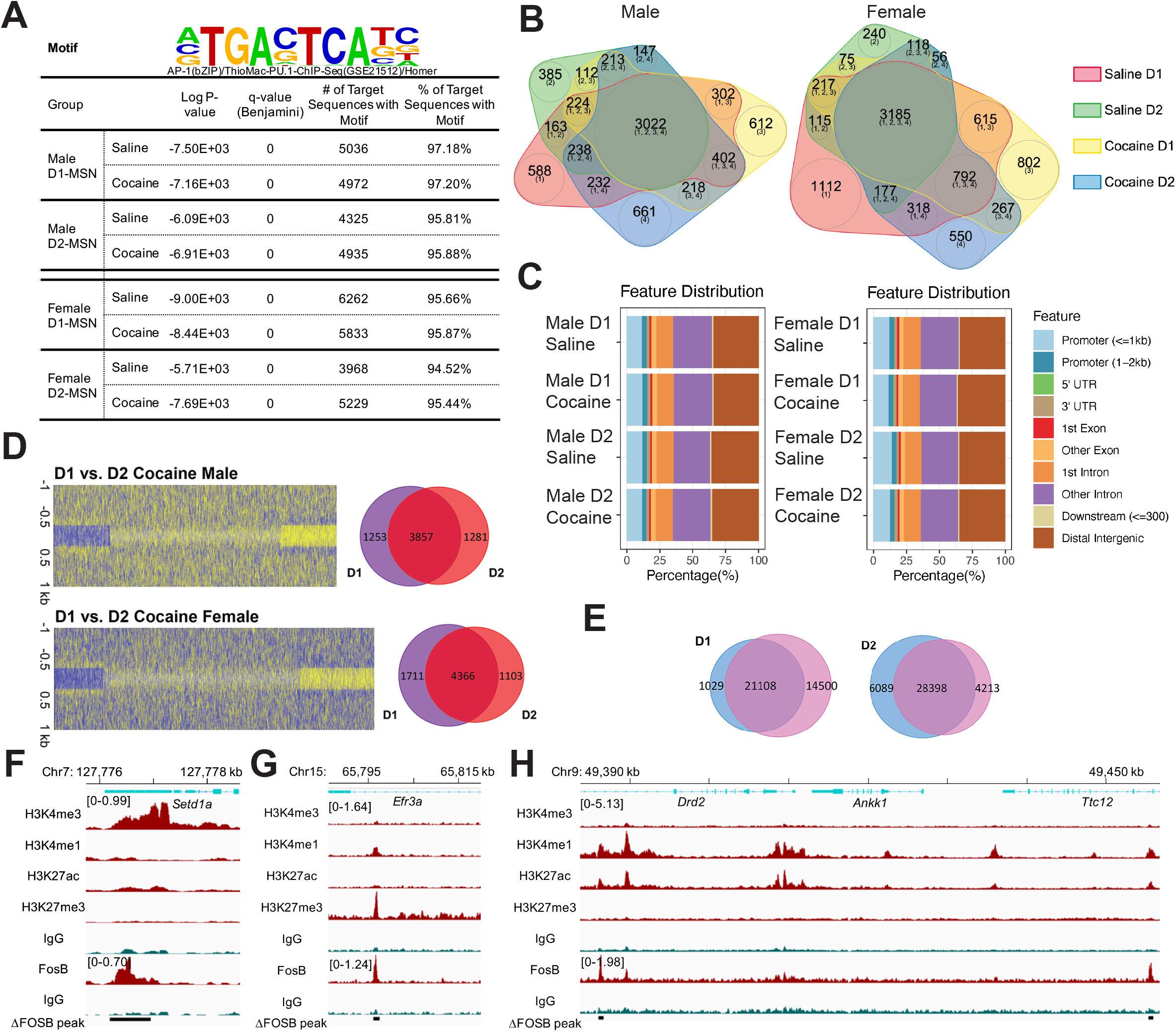
CUT&RUN-seq for ΔFOSB in NAc D1 MSNs and D2 MSNs after chronic cocaine exposure. (A) Motif analysis of called ΔFOSB peaks. (B) Venn diagram analysis of called ΔFOSB peaks among saline- and cocaine-exposed male and female mice. (C) Genomic distribution of ΔFOSB peaks in NAc D1 and D2 MSNs of male and female mice under saline and cocaine conditions. (D) Heatmaps comparing the effect of chronic cocaine on ΔFOSB peaks in D1 vs. D2 MSNs in male (upper) and female (lower) mice. (E) Venn diagrams comparing the effect of chronic cocaine on ΔFosB peaks in D1 and in D2 MSNs in male vs. female mice. (F) Example of ΔFOSB binding at a promoter region of *Setd1a* in the male D1 cocaine dataset. (G) Example of ΔFOSB binding at a poised enhancer region of *Efr3a* in the male D1 cocaine dataset. (H) Example of ΔFOSB binding at an active enhancer region in the *Drd2-Ankk1-Ttc12* locus in the male D2 cocaine dataset.

The greatest surprise from these datasets was the finding that roughly the same number of cocaine-induced changes in ΔFOSB binding, including induction of new peaks or increases or decreases in the amplitude of baseline peaks, occurred in D2 MSNs as in D1 MSNs (Fig. 2D), despite the overwhelming evidence that total nuclear levels of ΔFOSB are increased by cocaine in D1 MSNs only^5^. This observation held true in both sexes. These findings indicate that the role played by ΔFOSB in mediating aspects of cocaine action is not restricted to D1 MSNs and is broader than previously hypothesized.

It was also noteworthy that there was broad convergence in ΔFOSB binding sites between male vs. female mice, both in D1 and in D2 NAc MSNs. This is shown by Venn diagrams in Fig. 2E. This is striking contrast to the very small (<10%) convergence between males and females in the RNAs whose expression levels are altered in D1 and in D2 MSNs upon induction of the endogenous *Fosb* gene. See Discussion for consideration of these paradoxical findings.

### ΔFOSB binding sites are enriched within active enhancer regions

Historically, transcription factors were proposed to regulate gene expression by initiating transcription at specific gene promoter regions. More recently, several transcription factors have been shown to modulate transcriptional activity through binding at distant enhancer regions^17^. The genomic distribution of enhancer regions remains incompletely characterized, but in numerous tissues functional enhancers have been identified in gene bodies as well as in intergenic regions^18-20^. Based on our finding that the large majority of ΔFOSB binding sites occur in intergenic or gene body regions, we determined whether a subset of these ΔFOSB peaks are associated with enhancers. We accomplished this goal by performing cell-type-specific CUT&RUN-seq for several histone modifications that are linked with promoter or enhancer regions. Studies of enhancers in cultured neurons or non-neural tissues have found that they can be categorized into three groups: silenced, poised, and active enhancers, with specific histone modifications, H3K27me3 (histone H3 Lys27 trimethylation), H3K27me3/H3K4me1, and H3K27ac (acetylation)/H3K4me1, serving as indicators of these three groups, respectively^16^.

We identified tens of thousands of regions with these various histone modifications in D1 and D2 MSNs of the NAc. Through Venn diagram analyses, we found that all 8 experimental groups—cell type (D1 vs. D2 MSNs) x treatment (cocaine vs. saline) x sex (male vs. female)—show similar results: more than 70% of identified ΔFOSB peaks in both intergenic and gene body regions were surrounded by H3K27ac and H3K4me1 marks, a combination that marks active enhancers, with only ∼10% of ΔFOSB peaks displaying neither modification (Fig. 3A & S3A). The same degree of enrichment of ΔFOSB peaks at active enhancers in intergenic and gene body regions, shown for male D1 MSNs, was seen for male D2 MSNs and for both cell populations in the female NAc (Fig. S3A).

**Figure 3.**
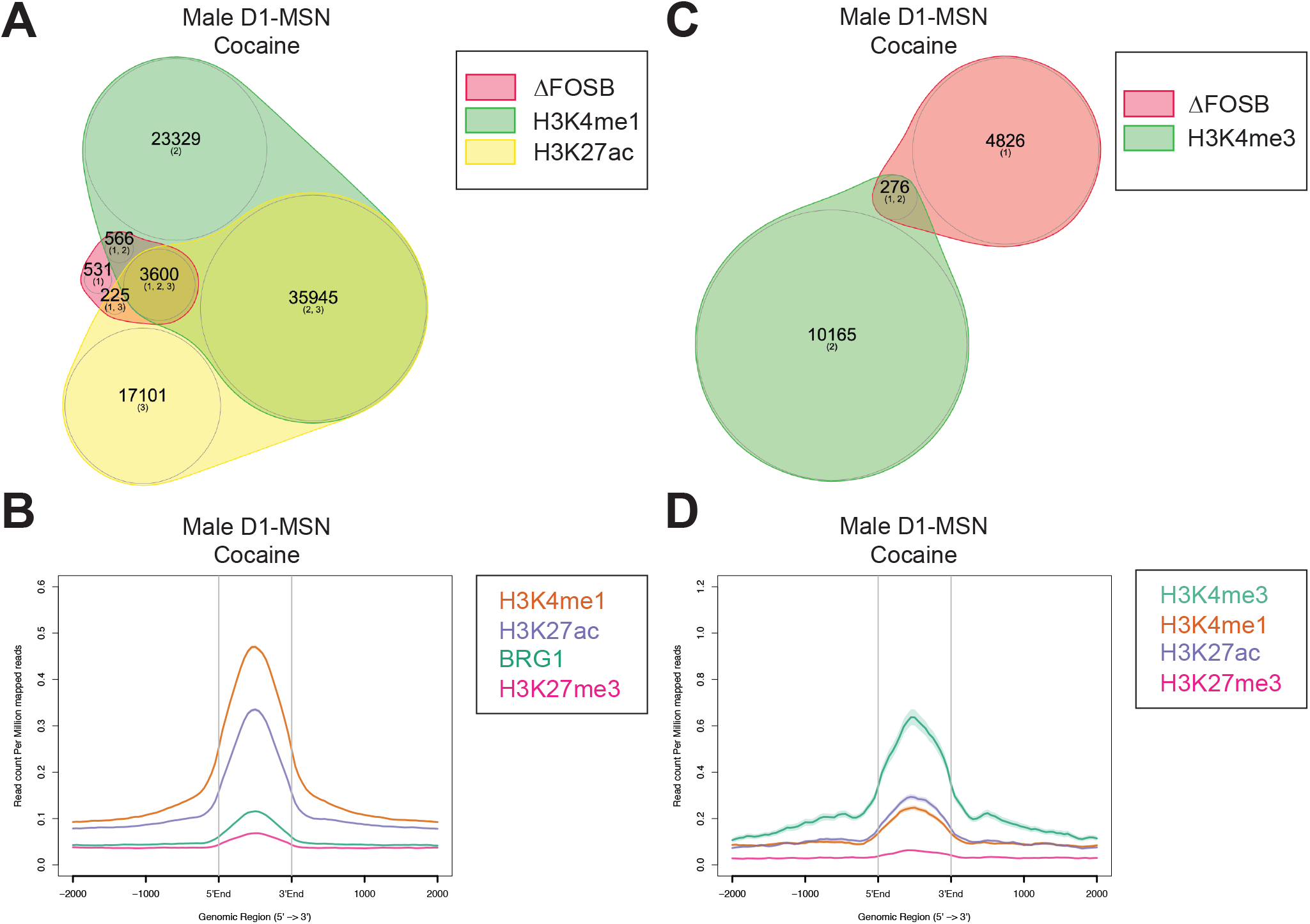
A majority (∼70%) of ΔFOSB binding sites overlap with BRG1 and with histone marks indicative of active enhancers. (A) Venn diagram analysis of called H3K4me1, H3K27ac, and ΔFOSB peaks in male D1 MSNs of NAc from cocaine mice. (B) Coverage of BRG1 and histone marks at called ΔFOSB+ / H3K4me1+ / H3K27ac+ peaks in the male D1 MSN cocaine group. (C) Venn diagram analysis of called H3K4me3 and ΔFOSB peaks in the male D1 MSN cocaine group. (D) Coverage of histone marks at called ΔFOSB+ / H3K4me3+ peaks in the male D1 MSN cocaine group.

The chromatin remodeling complex, SWI/SNF, controls enhancer activity by establishing the H3K27ac modification^22^, and we have reported previously that ΔFOSB forms a complex with BRG1 (the core subunit of the SWI/SNF complex) and the histone acetyltransferase, CBP/p300, which serves as a transcriptional coactivator, at a small number of putative candidate gene targets^23,24^. To examine whether the interaction between ΔFOSB and BRG1 is related to the enrichment of H3K27ac seen near ΔFOSB binding sites, we carried out BRG1 CUT&RUN-seq on NAc D1 and D2 MSNs and observed that sites of BRG1 enrichment overlapped with ΔFOSB sites that were also enriched for H3K4me1 and H3K27ac (Fig. 3B). The same general patterns were seen for D1 and D2 MSNs of both male and female mice (Fig. S3B).

Given that ∼15% of ΔFOSB peaks are located at promoter regions, and all prior studies of ΔFOSB binding to target genes focused on promoter regions (see Introduction), we performed CUT&RUN-seq for H3K4me3, a histone mark that is highly enriched at active gene promoters^21^. Approximately 5% of ΔFOSB peaks coincided with sites of H3K4me3 enrichment in male D1 MSNs (Fig. 3C) and D2 MSNs and both cell types in females (Fig. S3C). When examining ΔFOSB peaks located at promoters, only ∼50% of them overlapped with H3K4me3. This finding suggests that ΔFOSB binding to a gene’s promoter is not always associated with transcriptional activation, consistent with candidate gene and earlier gene expression array studies, which have suggested that ΔFOSB functions as a transcriptional repressor of certain genes^8,25-27^. Of note, ΔFOSB/H3K4me3 loci do not appreciably correspond to binding sites of histone marks related to enhancer activity (Fig. 3D & S3D) and, conversely, ΔFOSB peaks that overlap with enhancers show only weak H3K4me3 binding (data not shown), as would be expected based on studies of other tissues.

Examples of ΔFOSB binding peaks at individual gene targets, which illustrate the pattern of these various histone marks at a promoter region, at a poised enhancer, and at an active enhancer are shown in Fig. 2F, G, and H, respectively. Each of these genes, *Setd1a, Efr3a, Drd2*, and *Ttc12*, have not previously been implicated as ΔFOSB targets. Fig. S2C shows data for three genes that have been studied earlier as candidate target genes for ΔFOSB, namely, *Sirt2, Fos*, and *Camk2a*^10,26,28-30^. Our new CUT&RUN-seq data confirm these published findings. In contrast, several other candidate genes proposed to serve as targets for ΔFOSB—*Cdk5, Gria2*, and *Ehmt2*^31-33^—were not validated in our datasets. Possible explanations for these surprising findings are included in the Discussion.

### Motifs of homeobox and T-box transcription factors are adjacent to ΔFOSB-binding loci

Members of the FOS transcription factor family have been shown in cultured cells to recruit SWI/SNF complexes to maintain the accessibility of the nearby chromatin and to thereby act cooperatively with other transcription factors in a cell-type-specific manner to achieve coordinated regulation of gene expression^34^. Since BRG1 signal overlapped with ΔFOSB binding sites within active enhancer regions in our datasets, we performed a motif analysis on sequences 500 bp upstream and downstream of ΔFOSB binding loci to identify potential cooperative transcription factors for ΔFOSB in NAc D1 and D2 MSNs. We first located the consensus AP1 sequence in ΔFOSB binding sites and then searched for other nearby motifs. More than 100 motifs that were statistically significant (q-value < 0.05) genome-wide were then ranked based on the recurrence in target sequences. In addition to the AP1 motif, the top-ranked other motifs correspond to binding sites of several homeobox and T-box transcription factors (Fig. 4A & S4). There was no difference in these associations between D1 vs. D2 MSNs or between cocaine and saline conditions. This analysis for the first time implicates members of these two transcription factor families in ΔFOSB’s transcriptional machinery, with important implications for understanding pathological states in which ΔFOSB has been implicated.

**Figure 4.**
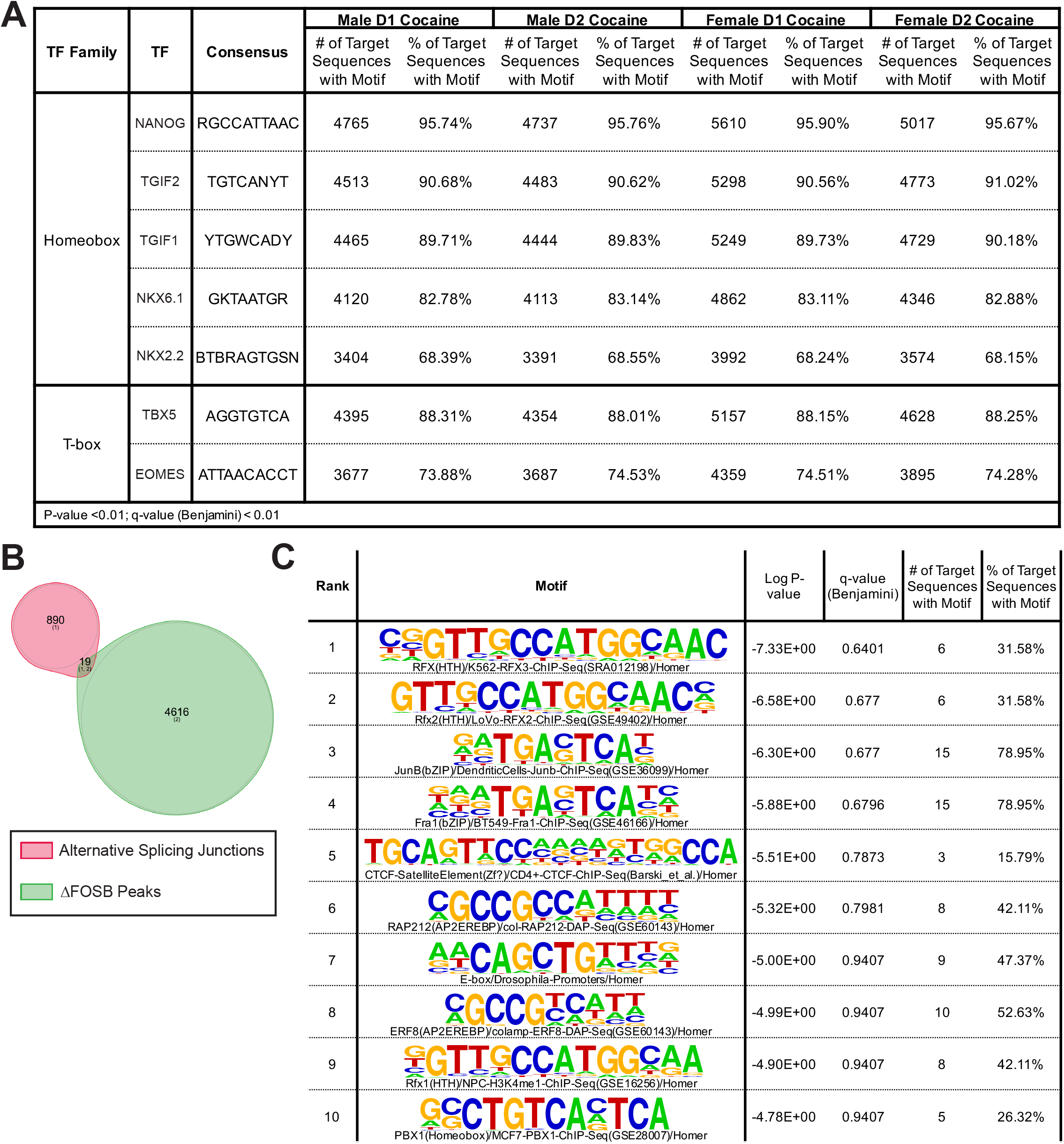
Loci with ΔFOSB binding sites contain motifs recognized by homeobox and T-box cription factors. (A) *In silico* motif analyses of loci with ΔFOSB-bound AP1 sites (± 500 bp from sites) in D1 and D2 MSNs of male and female cocaine groups. (B) Venn diagram analysis of native splicing junctions and ΔFOSB binding sites. (C) Motif analysis of alternative splicing ions with ΔFOSB peaks nearby.

### Lack of evidence for a role of ΔFOSB in regulating alternative splicing

Studies have shown that alternative splicing can be determined not only by the splicing machinery, but also by transcription factors and chromatin structure^35,36^. To examine whether ΔFOSB binding sites within gene bodies are related to alternative splicing, we compared the locations of ΔFOSB peaks to the alternative-splicing junctions identified in NAc under cocaine and saline self-administration conditions in a recent study^37^. Nine hundred and nine junctions represented differential alternative splicing events (p-value < 0.05 and Δ percentage spliced in > 0.1), but only 19 of them have detectable ΔFOSB binding sites within 1,000 bp upstream or downstream of their splice donor or acceptor sites (Fig. 4B-C). These 19 sites account for only 2% (19/909) of splicing junctions and 0.4% (19/4635) of all of the ΔFOSB binding sites identified in our D1 and D2 MSN datasets. This analysis indicates that the vast majority of ΔFOSB binding sites within gene body regions in the NAc do not overlap with alternative splicing junctions.

## Discussion

The induction of ΔFOSB in NAc has been demonstrated in humans with substance use disorders as well as in rodents after chronic exposure to cocaine or other drugs of abuse^1,2^. In addition, viral or genetic bidirectional manipulation of ΔFOSB activity in rodent NAc neurons controls the development of addiction-related behaviors^1,2,27^. However, it has been challenging to identify the direct genomic targets of ΔFOSB, beyond studies of a small number of candidate genes, due to technical limitations of using ChIP-seq to interrogate transcription factors in micro-dissected brain tissue. Here, by use of the newly developed CUT&RUN-seq procedure, which is far more sensitive and specific compared to previous ChIP-seq approaches, we have revealed, first, the whole-genome landscape of ΔFOSB binding sites selectively in NAc D1 and D2 MSNs at baseline and after chronic cocaine exposure. Second, we show that ∼85% of ΔFOSB binding sites are outside of known promoter regions and located within either distal intergenic regions or gene bodies. Third, a large majority of ΔFOSB peaks in intergenic regions and gene bodies are surrounded by histone marks indicative of active enhancers. Finally, our analyses suggest that ΔFOSB might function in a coordinated fashion with homeobox and T-box transcription factors to regulate gene targets in a cell-type-specific manner.

Identifying ΔFOSB’s target genes has been a persisting challenge for many years. Through manipulating the expression of ΔFOSB in NAc, including manipulating its expression in a cell-type-specific manner, several genes have been considered potential ΔFOSB targets due to changes in their expression levels upon overexpression or inhibition of ΔFOSB. However, we did not see significant binding of ΔFOSB around some of these candidate genes, for example, *Cdk5, Gria2*, and *Ehmt2*^31-33^, in our datasets, suggesting perhaps an indirect role of ΔFOSB in regulating these genes: namely, that ΔFOSB might control the expression of these genes through its regulation of intervening gene targets that encode transcriptional or chromatin regulatory proteins. An alternative hypothesis discussed further below is that ΔFOSB might indeed directly regulate these genes but through actions at distant enhancers that form loops with these loci. Characterization of the 3D structure of chromatin in D1 and in D2 MSNs under control and cocaine conditions will be needed to evaluate this possibility. By contrast, three other putative target genes implicated in previous work, *Fos, Sirt2*, and *Camk2a*^10,26,28-30^, were validated in our ΔFOSB CUT&RUN-seq dataset, confirming that they are bona fide direct targets of ΔFOSB. Our new datasets now provide for the first time a comprehensive map of all such genomic regions that are targeted by ΔFOSB in the NAc and can be used to identify and characterize numerous other genes whose expression levels are controlled in D1 and in D2 MSNs by this transcription factor at baseline and after a course of chronic cocaine exposure.

Another limitation in previous studies of ΔFOSB gene targets has been the assumption that ΔFOSB regulates genes primarily through binding at their promoter regions. All prior candidate gene studies examined ΔFOSB binding to gene promoters by use of ChIP-qPCR^1,26,28-30^, and the only prior effort to map ΔFOSB binding in NAc genome-wide used promoter arrays^10^. We now know that these efforts were in large part misplaced, given than only ∼15% of ΔFOSB peaks correspond to gene promoters. This novel insight underscores the importance of open-ended, unbiased approaches as utilized in the present study. Our finding that ΔFOSB binding is highly concentrated at active enhancers now fundamentally redirects the field’s efforts in identifying its target genes and understanding its mechanisms of transcriptional regulation. This will require more accurate and complete maps of the genes controlled by a given enhancer region, which must be determined for each cell type of interest. Indeed, our efforts benefit from a large repository of ChIP-seq and CUT&RUN-seq data for numerous histone modifications and chromatin-regulatory proteins in NAc, many in D1 and D2 MSNs selectively, which enable a comprehensive understanding of the regulation of gene expression through enhancer mechanisms in these two cell types^33-38^. Since enhancer function is also controlled by the 3D structure of chromatin, as noted above, which can bring a distant enhancer into close proximity to a gene promoter through chromosomal looping, it will be important to generate 3D maps of chromatin interactions, such as by use of HiC^39^, for D1 and D2 MSNs under control and cocaine conditions—maps which are not yet available.

The enrichment of ΔFOSB peaks in D1 and D2 MSNs at intergenic and gene body regions is unexpected since many transcription factors do bind predominantly at gene promoters. While this has been demonstrated mainly in cultured cells and peripheral tissues, it presumably holds true for brain as well. Consistent with this expectation, CUT&RUN-seq maps for another cocaine-regulated transcription factor, CREB (cAMP response element binding protein)^7^, in D1 and D2 MSNs shows >80% of CREB peaks at gene promoters (data not shown). Nevertheless, there are precedents for transcription factor binding sites being located within gene bodies, especially within introns as seen for ΔFOSB in the present study, a feature demonstrated for example for forkhead transcription factors and the tumor suppressor, p53.^40-42^ Moreover, there increasing evidence for active enhancers also being located within gene body regions,^18-20^ and even reports that an enhancer present within the body of one gene can influence the expression of a distinct gene.^43^ Our findings with ΔFOSB therefore are consistent with this rapidly evolving literature and point to important features of its transcriptional actions that set it apart from certain other neural transcription factors (e.g., CREB).

A third variable in studies of ΔFOSB gene targets is the use of a particular anti-FOSB antibody with proper IgG controls. This is exemplified by the three candidate gene targets, *Fos, Sirt2*, and *Camk2a*, mentioned earlier^10,26,28-30^. The anti-FOSB antibodies utilized for these previous ChIP-qPCR experiments were all different. Moreover, while these published studies focused on ΔFOSB binding at promoter regions only, we detected a significant ΔFOSB peak solely within the *Camk2a* gene body, while a strong signal at the promoter region was not called due to significant noise in the IgG control (Fig. S2). On the other hand, this promoter binding site was lost in the NAc of *Fosb-*null mice. Therefore, further work is needed to determine whether the ΔFOSB peak within the *Camk2a* gene promoter represents a functional ΔFOSB binding site or not. These considerations emphasize the importance of using control conditions to verify bona fide ΔFOSB binding, such as loss of signal in *Fosb*-null mice and absence of signal with an IgG control.

Multiple studies have shown that cellular or nuclear levels of ΔFOSB in NAc are induced after chronic cocaine exposure, an adaptation specific to D1 MSNs after either investigator- or self-administered drug^1,6,13^. It is therefore striking that we observed a comparable number of cocaine-induced changes in ΔFOSB binding in D2 MSNs as found for D1 MSNs. This observation underscores that the total nuclear level of a transcription factor is not in and of itself an accurate reflection of the genomic regulation mediated by that factor. Indeed, we found depletion of ΔFOSB binding at a subset of loci in response to chronic cocaine despite induction of the protein. These data emphasize the importance of obtaining the genome-wide maps for a transcription factor as accomplished in this study for ΔFOSB. FOS family proteins form active AP1 transcription factor complexes by heterodimerizing with a JUN family protein, and prior work confirmed the enrichment of ΔFOSB:JUND heterodimers in brain^44^. Interestingly, ΔFOSB has also been shown to form homodimers, uniquely among all other FOS proteins, and ΔFOSB:ΔFOSB homodimers might display distinct gene regulation patterns^45^. It will therefore be important in future studies to obtain genome-wide maps of JUND and perhaps other JUN family proteins to distinguish between loci of ΔFOSB heterodimer vs. homodimer binding. Moreover, the anti-FOSB antibody used in this study recognizes both full-length FOSB and ΔFOSB. We assume that all of the signals in our CUT&RUN-seq datasets reflect ΔFOSB only, since full-length FosB is not detected by Western blotting under the saline and cocaine conditions used in this^1,6,13^. Nevertheless, we cannot rule out the contribution of full-length FosB in the absence of an antibody that recognizes ΔFOSB specifically, a difficult undertaking since the full ΔFOSB protein sequence is contained within full-length FOSB.

ΔFOSB has long been considered a key player in the development of drug addiction, but it has also been implicated in numerous other brain functions and neuropsychiatric disorders, including stress susceptibility and resilience, learning and memory, Alzheimer’s disease, and epilepsy, among others^2,3,46-49^. Accordingly, there is interest in developing small molecule potentiators or inhibitors of ΔFOSB as possible imaging agents for diagnostic purposes or even as novel therapeutic approaches^2,50^. The present study expands our understanding of ΔFOSB’s molecular functions and will assist strategies to control ΔFOSB’s transcriptional machinery. Increasing evidence indicates that certain transcription factors function in cooperation with several other factors, with a major area of current research focused on delineating such cooperative partners. Indeed, we found that motifs of homeobox and T-box transcription factors are located ±500 bp from the large majority of all identified ΔFOSB binding sites. Further pinpointing the specific homeobox and T-box transcription factors that interact with ΔFOSB to regulate expression will shed light on ΔFOSB’s transcriptional actions as well as how to develop effective therapeutic approaches for drug addiction and other conditions. The homeobox transcription factors, PKNOX2 and CUX2, have been linked to substance dependence and excessive alcohol consumption, repectively^51,52^. Studies are now needed to investigate whether these factors underlie the cooperativity we deduced bioinformatically. Lastly, one of the cell-type-specific ΔFOSB binding sites that we identified in NAc D2 MSNs corresponds to the *Drd2-Ankk1-Ttc12* locus, which is a gene cluster associated with human substance use disorders^53^. Overall, the datasets generated in this study will therefore not only decipher ΔFOSB’s molecular functions, but also help fill the gaps between rodent models and findings derived from human genetics.

In summary, through combining genetic, molecular biological, and bioinformatic approaches, we elaborate the cell-type-specific binding sites for ΔFOSB genome-wide in D1 MSNs and D2 MSNs of the NAc at baseline and after chronic cocaine exposure. The large majority of ΔFOSB peaks identified correspond with histone marks that are indicative of active enhancers. It will now be possible to leverage these new datasets, combined with other genome-wide maps such as for 3D chromatin structure, to delineate the full range of target genes whose expression levels are altered through ΔFOSB-dependent mechanisms in MSN subtypes. Our *in silico* results highlight the potential of homeobox and T-box transcription factors in coordinating with ΔFOSB to control target gene expression, another fruitful focus for future investigations. This study thereby provides a critical step forward in understanding the role of ΔFOSB in the development of drug addiction and other neuropsychiatric disorders as well as ultimately mining these discoveries for therapeutic innovations.

## Acknowledgements

This work was supposed by grants from the National Institute on Drug Abuse (P01DA047233 and R01DA007359) to EJN.

## Methods

### CONTACT FOR REAGENT AND RESOURCE SHARING

Further information and requests for resources and reagents should be directed to, and will be fulfilled by the corresponding author, Dr. Eric J. Nestler.

### ANIMAL MODELS

#### Mouse lines

Mice were housed in an AAALAS-certified facility at the Icahn School of Medicine at Mount Sinai and all animal protocols were approved by the Institutional Animal Care and Use Committee. Mice were maintained on a 12-h light/dark cycle (7am ON, 7pm OFF) and given access to food and water ad libitum. Intravenous self-administration was maintained on a reverse 12-h light/dark cycle (7am OFF, 7pm ON). The following lines of mutant mice were used: *Drd1-Cre* (Tg(Drd1-cre)FK150Gsat/Mmucd, GENSAT), *Drd2-Cre* (Tg(Drd2-cre)ER44Gsat/Mmucd, GENSAT), *ROSA-Sun1/sfGFP* (B6;129-*Gt(ROSA)26Sor*^*tm5(CAG-Sun1/sfGFP)Nat*^/J, JAX 021039), *Sox2-Cre* (B6.Cg-*Edil3*^*Tg(Sox2-cre)1Amc*^/J, JAX 008454), *Fosb-flox*^15^ and wild-type C57BL/6J (JAX 000664). Genomic PCR was used to distinguish wild-type *Fosb, Fosb*-floxed and *Fosb*-null alleles with the following PCR genotyping primers (wild-type, 359 bp; *Fosb*-floxed, 397 bp; *Fosb*-null: 175 bp): Fosb_CKO_F: 5’-TGCCATAACTAGTCCTTGGAAGTTC-3’ Fosb_flox_F: 5’-GCTGAAGGAGATGGGTAACAGAAC-3’ Control_R: 5’-GCAAGAACTCCAAGCCTGGT-3’

#### Cocaine injections

Eight-to-ten-week-old mice received daily intraperitoneal (IP) injections of saline (0.9% NaCl) or cocaine (20 mg/kg) dissolved in saline. Tissues were collected 24 hr after the last injection.

#### Cocaine self-administration

Mice maintained on mild, overnight food restriction were first trained to lever press for a food reward in operant conditioning chambers (Med Associates; Fairfax, VT). Two levers were presented to mice at the beginning of the session, and responding on the active lever led to the delivery of a chocolate pellet (Bio-Serv; Flemington, NJ) on a fixed-ratio 1 (FR1) schedule of reinforcement. Responding on the inactive lever had no consequence. Sessions lasted for 1 h or until 30 reward pellets had been earned. Mice were trained until acquisition criteria for lever pressing had been met (30 pellets earned in two consecutive sessions with <10 inactive lever responses; 4 sessions on average required).

Following acquisition of responding for food, mice underwent surgical procedures under inhaled isoflurane anesthesia (2%) to implant a pre-constructed intravenous catheter (Strategic Applications Incorporated; Lake Villa, IL; SBD-05C) in the right jugular vein. The catheter cannula exited the skin from the animal’s mid-back, and catheter tubing (0.013 inch inner diameter) was threaded subcutaneously over the right shoulder and inserted 1.3 cm into the jugular vein. Ketoprofen (5 mg/kg) was injected subcutaneously following surgery as post-operative analgesia. Mice were allowed to recover for 4 days, during which catheters were flushed once daily with 0.03 ml heparinized saline (30U) containing ampicillin antibiotic (5 mg/ml) to forestall infection.

Following recovery, mice started cocaine or saline self-administration in the operant conditioning chambers, wherein responding on the active lever, which previously delivered food, now delivered an intravenous infusion of cocaine (0.5 mg/kg/infusion) on a FR1 schedule of reinforcement in 2 hr daily sessions over 10 days. Infusions were signaled by the activation of a cue light located above the active lever, and were followed by a 20 sec timeout during which the cue light remained illuminated, and levers remained extended. Responding on the active lever during timeout was recorded but had no consequence. Throughout the session, responding on the inactive lever had no consequence. Catheters were flushed with heparinized saline (30U) before and after self-administration sessions. Twelve hr following the last self-administration session, catheter patency was tested. During self-administration procedures mice were maintained on ad libitum food access. Mice were flushed with 0.03 ml of 3 mg/ml methohexital sodium dissolved in saline, followed immediately by 0.03 ml of saline. Patency was confirmed by loss of muscle tone within 3 sec following infusion. Mice lacking catheter patency were excluded. Tissues were collected 12 after the patency test (i.e., 24 hr after the final self-administration session).

### TISSUE COLLECTION

Mouse brains were dissected and place in ice-cold PBS on ice after cervical dislocation. The brains were placed into an ice-cold brain slicer and sliced into 1 mm thick coronal sections with razor blades. NAc and dorsal striatum from both hemispheres were harvested with 14-gauge syringe needles from the 1 mm thick brain slice and punches were flash-frozen and stored at −80°C for further applications.

### CUT&RUN-SEQ

#### Nuclei preparation

Nuclei were isolated from frozen tissue. Punches were homogenized with Dounce homogenizers in 4 ml lysis buffer (320 mM sucrose, 5mM CaCl_2_, 0.1 mM EDTA, 10 mM Tris-HCl, pH 8.0, 1 mM DTT, 0.1% Triton X-100, 1.5 mM spermidine, and protease inhibitor), with loose pestle 30 times and tight pestle 30 times, and filtered with a 40 μm strainer. A discontinuous sucrose layer (1.8 M sucrose, 10 mM Tris-HCl, pH 8.0, 1 mM DTT, 1.5 mM spermidine, and protease inhibitor) was placed beneath the lysate and samples were centrifuged at 71,124 RCF for 1 h at 4°C, Nuclei in the pellet were resuspended on ice in 1 ml wash buffer (20 mM HEPES-NaOH, pH 7.5, 150 mM NaCl, 0.5 mM spermidine, 0.1% Triton X-100, 0.1% Tween-20, 0.1% BSA, and protease inhibitor).

#### INTACT nuclear isolation

Nuclei samples were pre-cleared with Dynabeads™ Protein G (10003, ThermoFisher) for 30 min at 4°C and then with GFP-Trap Magnetic Particles M-270 (gtd-20, ChromoTek GmbH) for 1 h at 4°C. Note that both beads were pre-washed twice with 1 ml wash buffer. The nuclei-bound beads were resuspended in 1 ml antibody buffer (wash buffer with 2 mM EDTA), inverted, and reclaimed at the magnet; this wash pattern was repeated 8 times. The nuclei-bound beads were then resuspended in 10 ml antibody buffer, inverted, and reclaimed at the magnet; this wash pattern was repeated 2 times. Finally, the beads were resuspended in 1 ml wash buffer, inverted, and reclaimed at the magnet; this final wash pattern was repeated 2 times.

### CUT&RUN

Bulk nuclei or INTACT-isolated nuclei were incubated with BioMag®Plus Concanavalin A (BP531, Bang Laboratories) for 10 min at room temperature. Note that the beads were activated twice with 1.5 ml binding buffer (20 mM HEPES-KOH, pH 7.9, 10 mM KCl, 1 mM CaCl_2_, and 1 mM MnCl_2_). The nuclei-bound beads were reclaimed at the magnet and incubated in antibody buffer containing specific primary antibody at 4°C overnight. The following antibodies were utilized in this study: anti-FOSB (2251, Cell Signaling, 1:50 dilution), anti-H3K27Ac (ab4729, Abcam, 1:50 dilution), anti-H3K4me1 (ab8895, Abcam, 1:50 dilution), anti-H3K4me3 (13-0041, EpiCypher, 1:50 dilution), anti-H3K27me3 (9733, Cell Signaling, 1:50 dilution), anti-BRG1 (13-2002, EpiCypher, 1:50 dilution), and non-immune rabbit IgG (2729, Cell Signaling, 1:100 dilution). The nuclei-bound beads were washed with antibody buffer and incubated with antibody buffer containing guinea pig anti-rabbit secondary antibody (ABIN101961, antibodies-online Inc., 1:100 dilution) at 4°C for 1 h. Afterward, nuclei-bound beads were washed with antibody buffer and incubated with antibody buffer containing pAG-MNase (15-1116, EpiCypher) at 4°C for 1 h. The beads were then washed with wash buffer twice and low-salt buffer (20 mM HEPES-NaOH, pH 7.5, 0.5 mM spermidine, 0.1% Triton X-100, 0.1% Tween-20, and protease inhibitor) once before being incubated in calcium incubation buffer (3.5 mM HEPES-NaOH, pH 7.5, 10 mM CaCl_2_, 0.1% Triton X-100, and 0.1% Tween-20) for 5 min on ice. The beads were reclaimed at the magnet and the reaction was stopped by addition of EGTA-stop buffer (170 mM NaCl, 20 mM EGTA, 0.1% Triton X-100, 0.1% Tween-20, 25 μg/ml RNase, and 20 μg/ml glycogen). DNA fragments were eluted by incubating the samples at 37°C for 30 min with 500 rpm mixing. The samples were centrifuged at 16,000 RCF for 5 min at 4°C and the supernatant was reserved for DNA clean-up using NucleoSpin® Gel and PCR Clean-Up (740609, Takara) based on manufacture instructions or ethanol precipitation.

#### Library preparation

Sequencing libraries were prepared using the NEBNext® Ultra™ II DNA Library Prep Kit for Illumina® (E7645, New England BioLabs) with NEBNext® Multiplex Oligos for Illumina® (Dual Index Primers Set 1) (E7600S, New England BioLabs) based on manufacture instructions. Libraries were submitted for sequencing by GENEWIZ on Illumina HiSeq 4000 or NovaSeq machine with a 2 × 150-bp paired-end read configuration to a minimum depth of 25 million reads.

### DATA ANALYSIS

#### Reference genome alignment

The NGS-Data-Charmer pipeline (Github commit date: June 18, 2020) was used for preprocessing, QC, and alignment of the dataset (https://github.com/shenlab-sinai/NGS-Data-Charmer). In brief, trim-galore (v0.6.5) was used in an initial adaptor trimming step. Any reads for which no adaptor sequence was removed during the initial trimming step underwent a secondary trimming step, in which cutadapt^54^ (v2.10) was used to remove 6 basepairs from the end of the reads (cutadapt -u -6). Read alignment was performed with Bowtie2^55^ (v2.4.1) (--dovetail --phred33) to the mm10 genome. Duplicated read pairs were subsequently removed using the samtools^56^ (v1.10) ‘rmdup’ module. For visualization purposes, TDF files were generated using IGVtools^57-59^ (v2.5.3) (igvtools count), and bigwig files were generated using Deeptools^49^ (v3.5.0) ‘bamCoverage’ module (bamCoverage --binSize 10 --normalizeUsing RPKM). For generating combined bigwig files, bam files were merged using samtools ‘merge’ module and corresponding index files were created using samtools ‘index’ module; bigwig files were then generated using Deeptools ‘bamCoverage’ module (bamCoverage --extendReads --ignoreDuplicates --centerReads --normalizeUsing CPM --binSize 10).

#### Peak calling

For each biological replicate and corresponding IgG control, peaks were called with MACS2^61^ (v2.2.6) (macs2 callpeak -f BAMPE --pvalue 0.05 --keep-dup) and filtered through Irreproducible Discovery Rate (IDR)^62^ (v2.0.4.2) using 1% IDR threshold for bulk NAc datasets and 5% IDR threshold for NAc D1 and D2 MSN datasets. IDR output for each group was combined with Bedtools^63^ (v2.29.2) ‘merge’ module and the output containing high confidence peaks were utilized for further analysis. ChIPseeker^53^ (v1.22.1) was used to annotate peaks in R (v3.6.1). Venn diagram analyses were carried out using HOMER (v4.9) ‘mergePeaks’ module (mergePeaks -d given). Differential peaks were identified using DiffBind package (v2.10.0)^65^ in R (v3.5.3).

#### Motif analysis

Motifs in the peaks were identified using HOMER software^66^ ‘findMotifGenome.pl’ module with scrambled sequence as control (findMotifsGenome.pl -size given). Motif tables analyzed were subsets of knownResults.

### RNA EXTRACTION AND QPCR

Brain tissue was homogenized in Trizol reagent (ThermoFisher) and RNA was extracted with Direct-zol RNA Microprep Kit (R2060, Zymo Research) based on manufacture instructions. Reverse transcription was performed with iScript cDNA Synthesis Kit (1708890, Bio-Rad) and qPCR was carried out through SYBR Green Master Mix (A25741, ThermoFisher) and the primer sets below:

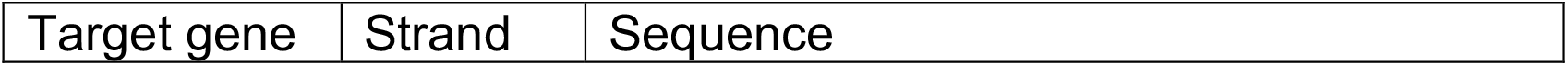
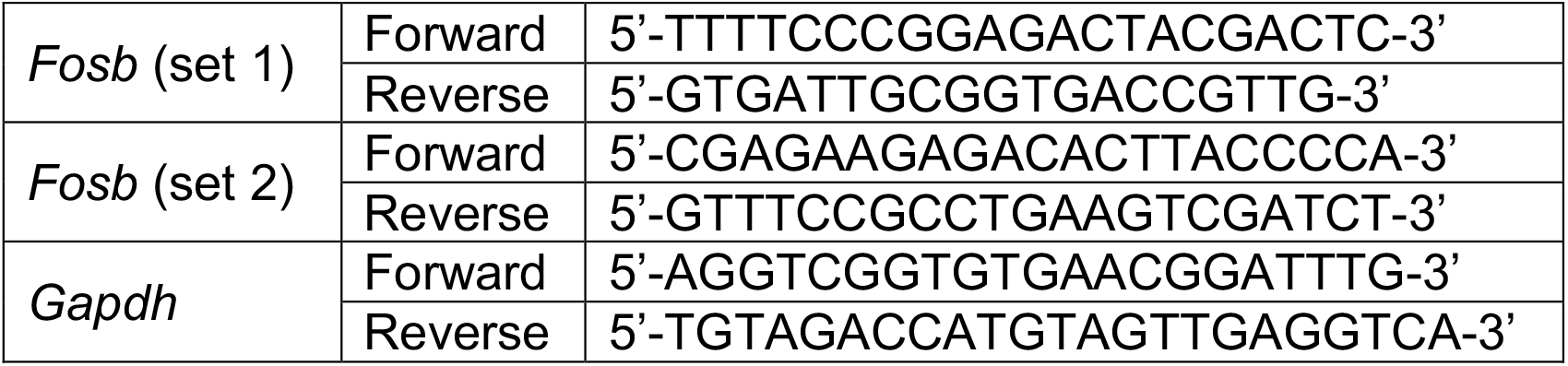

### PROTEIN EXTRACTION AND WESTERN BLOTTING

Brain tissue was homogenized in N-PER Neuronal Protein Extraction Reagent (P187792, ThermoFisher) with Halt™ Protease and Phosphatase Inhibitor Cocktail, EDTA-free (P178441, ThermoFisher) and protein samples were denatured in 4xLaemmli Sample Buffer (1610747, Bio-Rad) containing 2-mercaptoethanol. Western blotting was performed using the following antibodies: anti-vinculin (MCA465GA, Bio-Rad; 1:1,000 dilution), anti-FOSB (2251, Cell Signaling; 1:500 dilution), goat anti-rabbit antibody conjugated with HRP (G21234, ThermoFisher; 1:5,000 dilution), and goat anti-mouse antibody conjugated with HRP (G21040, ThermoFisher; 1:5,000 dilution).

### DATA VISUALIZATION

*Motif analysis* (Fig. 1A, S1A-B, 2A, S2A-B, and 4C): peaks were processed using HOMER^66^ ‘findMotifGenome.pl’ module.

*Venn diagram analysis* (Fig. 1B, 2B, 3A, 3C, S3A, S3C and 4B): output Venn diagram number were utilized to create Venn diagram plots using nVenn^67^ (v0.2.3), rsvg (v2.1.2), and grImport2 (v0.2-0) packages in R (v3.6.1).

Feature distribution plots (Fig. 1C and 2C): peaks were annotated by ChIPseeker package^64^ ‘annotatePeak’ function and visualized with ‘plotAnnoBar’ function in R.

Coverage heatmaps (Fig. 1E): Read density across 2 kb upstream and downstream of the peaks was obtained from combined bigwig files using Deeptools^60^ ‘computeMatrix’ module (computeMatrix reference-point --referencePoint center -a 2000 -b 2000) and visualized with ‘plotHeatmap’ module.

Coverage profile plots (Fig. 3B, 3D, S3B, and S3D): Read density across 2 kb upstream and downstream of the peaks was obtained from aligned bam files and visualized through ngs.plot^68^ (v2.61) (ngs.plot.r -G mm10 -L 2000) in R (v3.0.3).

Genome browser tracks (Fig. 2F-H, and S2C): combined bigwig files and peak files were inported into IGV^57-59^ (v2.9.4) with group autoscaling setting.

## Data and Software Availability

The CUT&RUN-seq data generated for this study have been deposited onto GEO under accession number GSE197668. RNA-seq and the results of alternative splicing analyses were deposited in early study^69^ under GEO accession number GSE141520.

## Author Contributions

S.-Y.Y. and E.J.N. conceived and designed the experiments. S.-Y.Y. and C.K.L. optimized and performed the experiments. S.-Y.Y., M.E. and L.S. performed data analysis. M.E. and L.S. provided software and general computational support. C.J.B., A.M.-T., and R.F. generated the self-administration mouse model. K.B. and C.M. assisted in generating the mouse models. S.-J. X., S.Z., and E.A.H. analyzed alternative splicing data. S.-Y.Y. and E.J.N. drafted the original manuscript, and all authors assisted in editing the manuscript.

---

**Figure S1.**
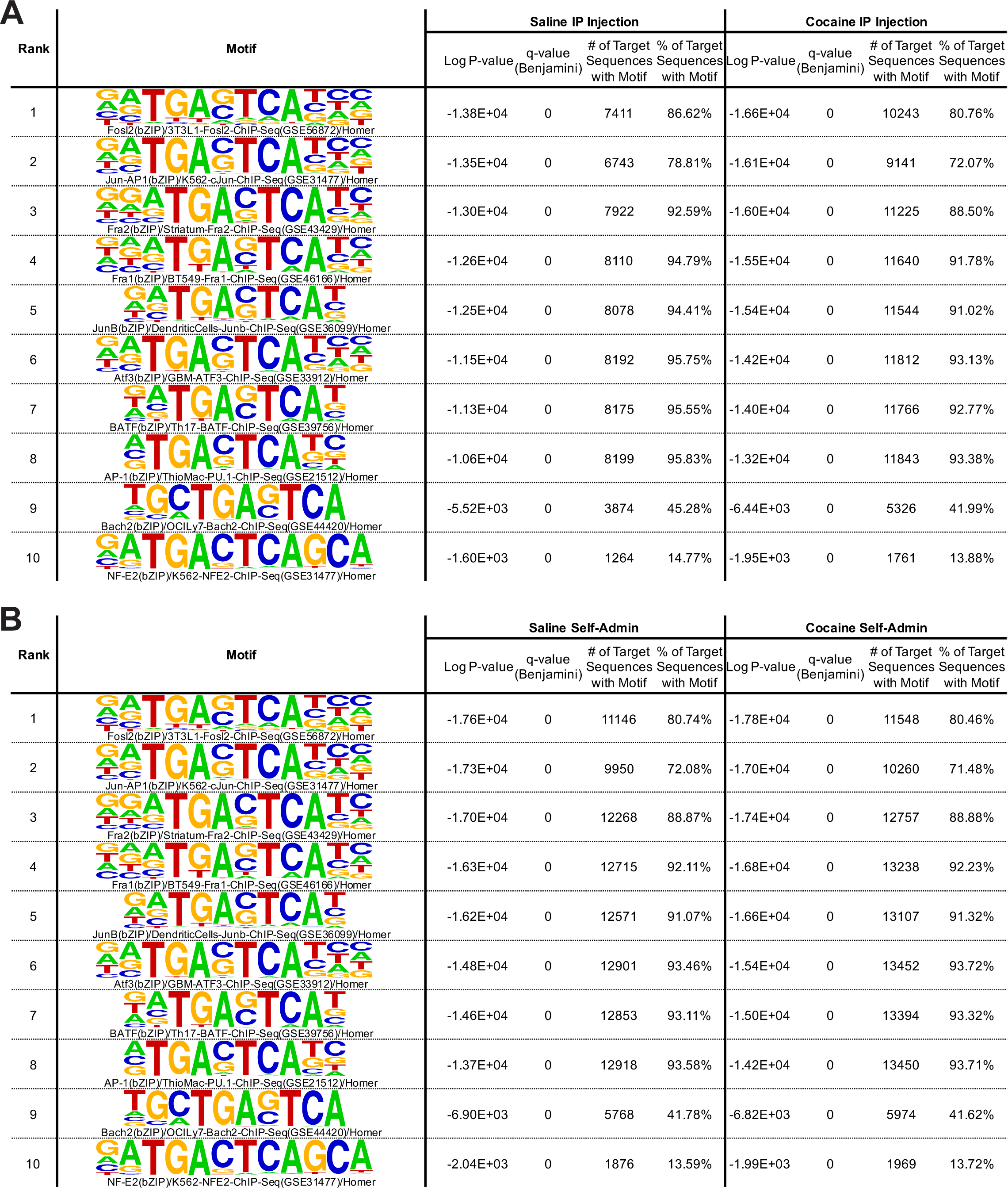
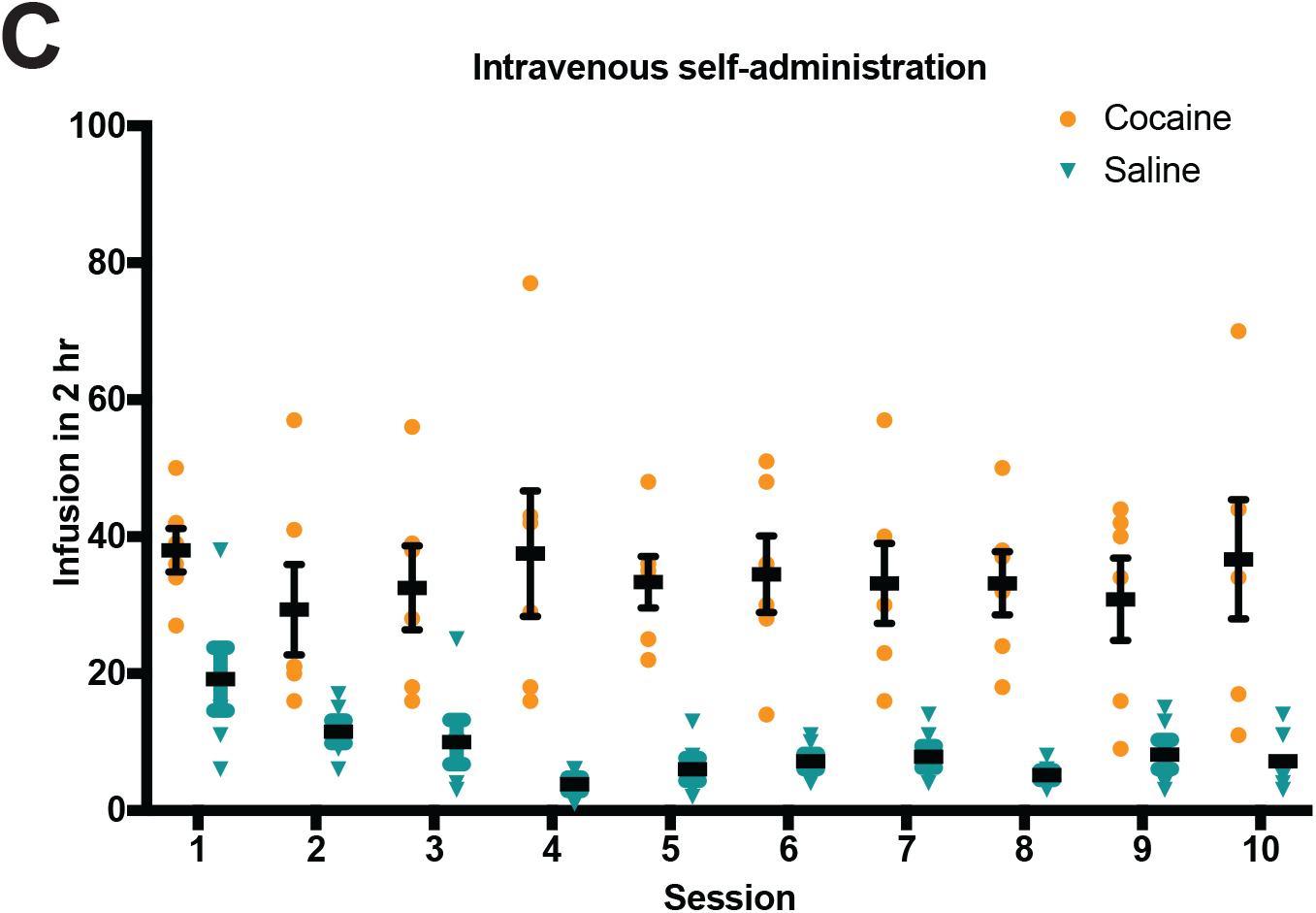
Top 10 motifs in the motif analyses of called ΔFOSB peaks in bulk NAc from male mice with cocaine (A) IP injections or (B) self-administration. (C) Infusion numbers (mean ± SE) of cocine/saline self-administering mice during the 10 daily 2-hr sessions.

**Figure S2.**
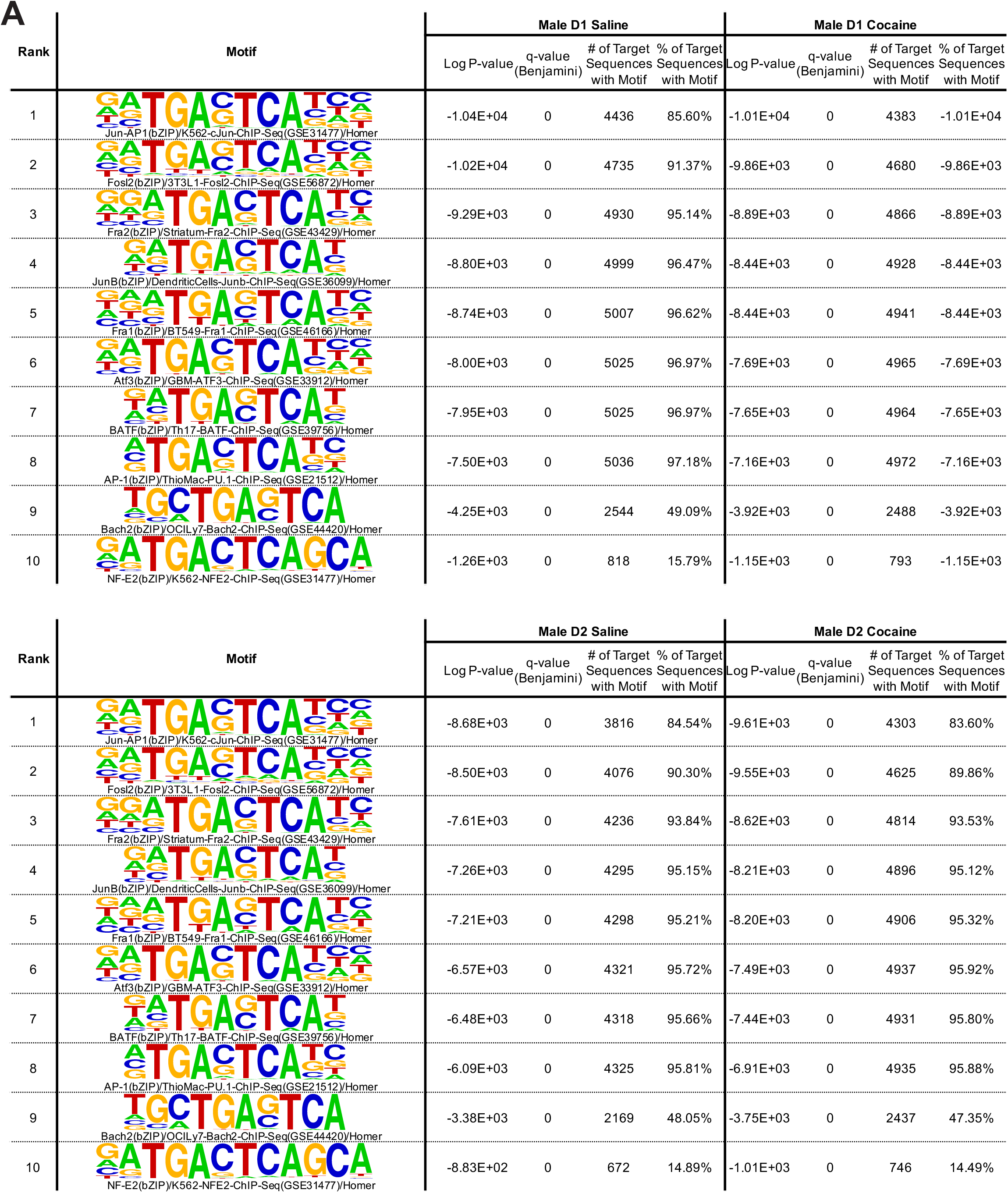
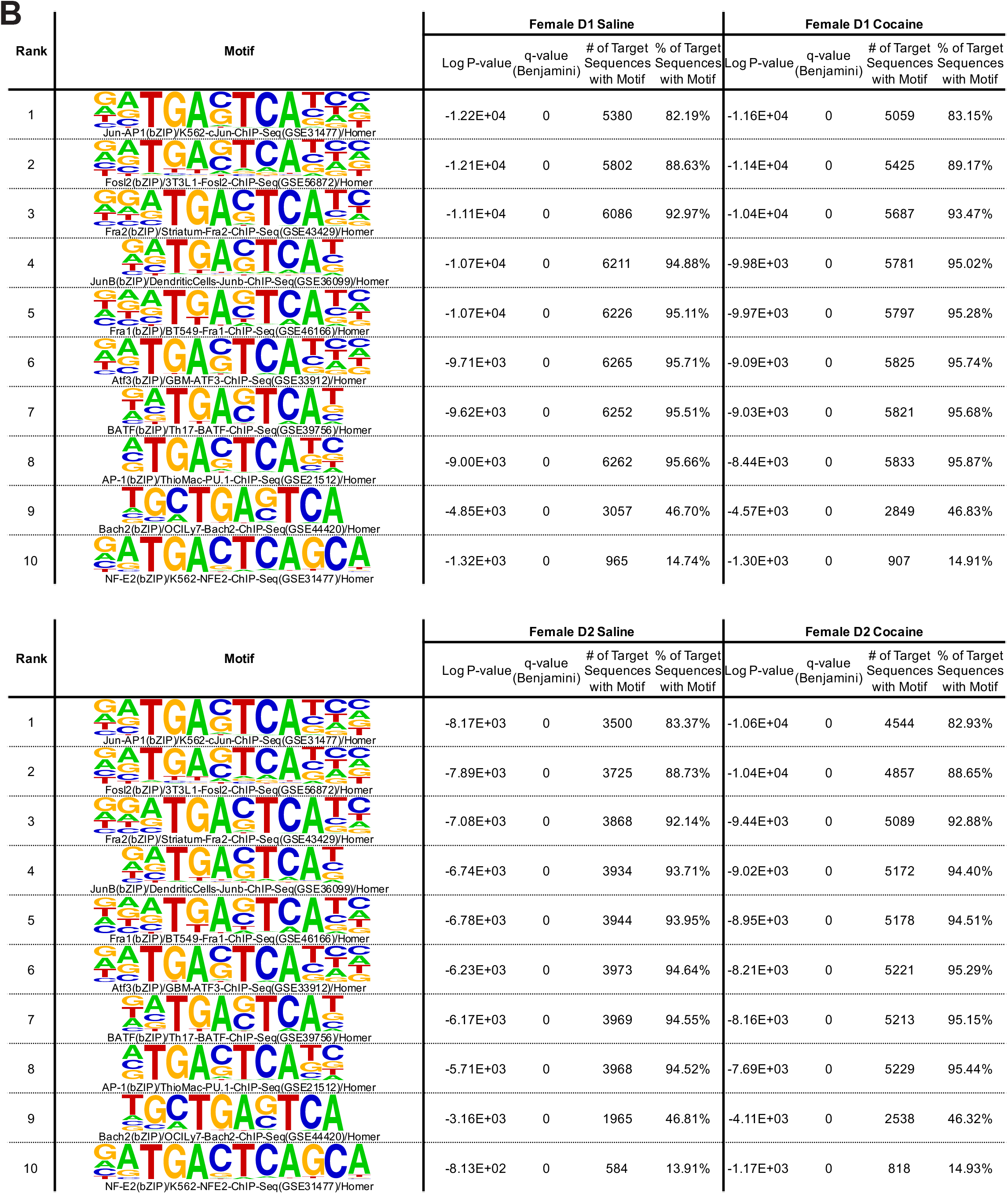
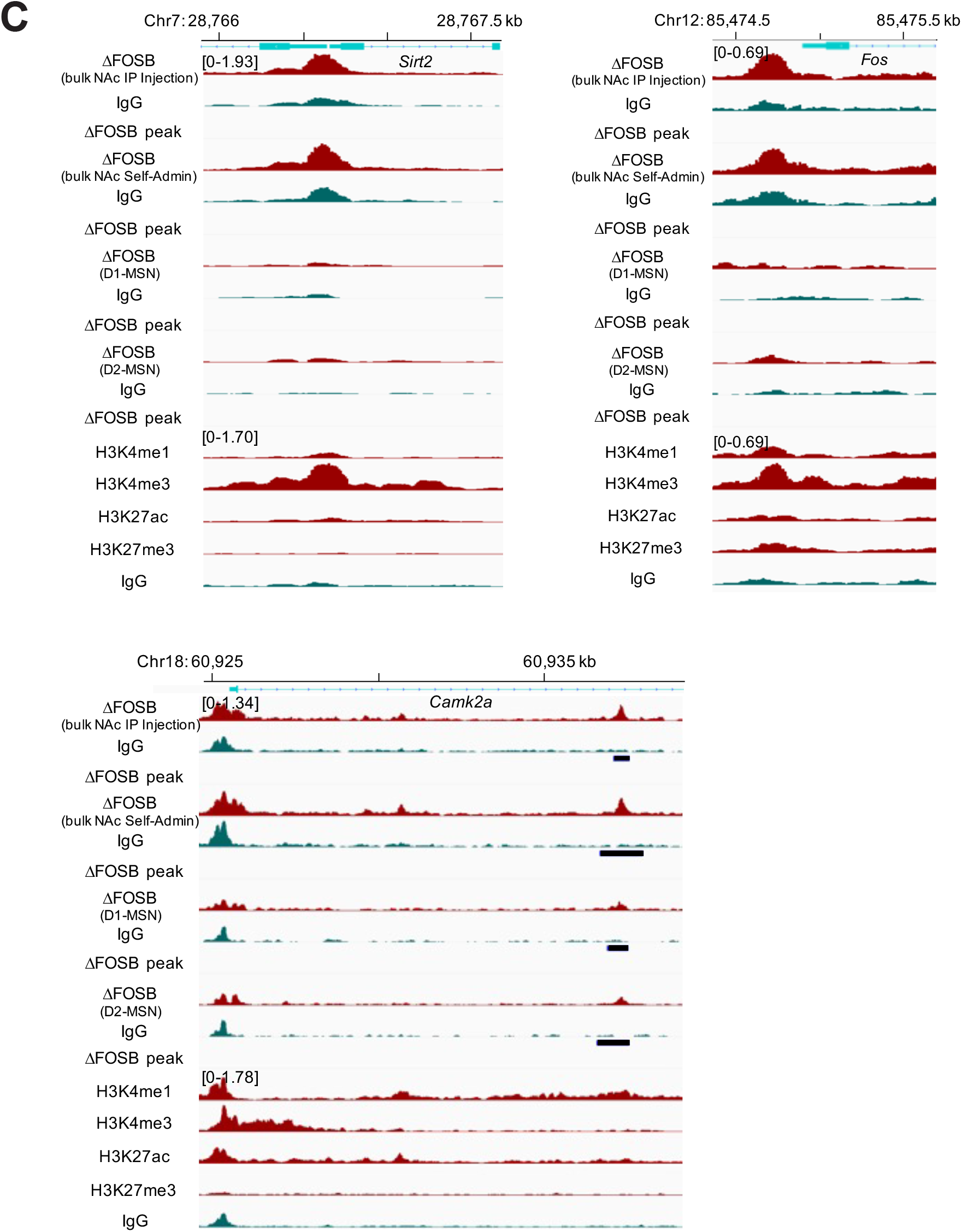
Top 10 motifs in the motif analyses of called ΔFOSB peaks in NAc D1 and D2 MSNs from (A) male and (B) female mice with cocaine or saline IP injections. (C) Tracks of ΔFOSB and histone mark CUT&RUN-seq at the regions of previously examined candidate target genes of ΔFOSB. Histone mark tracks were from male D1 MSN cocaine dataset.

**Figure S3.**
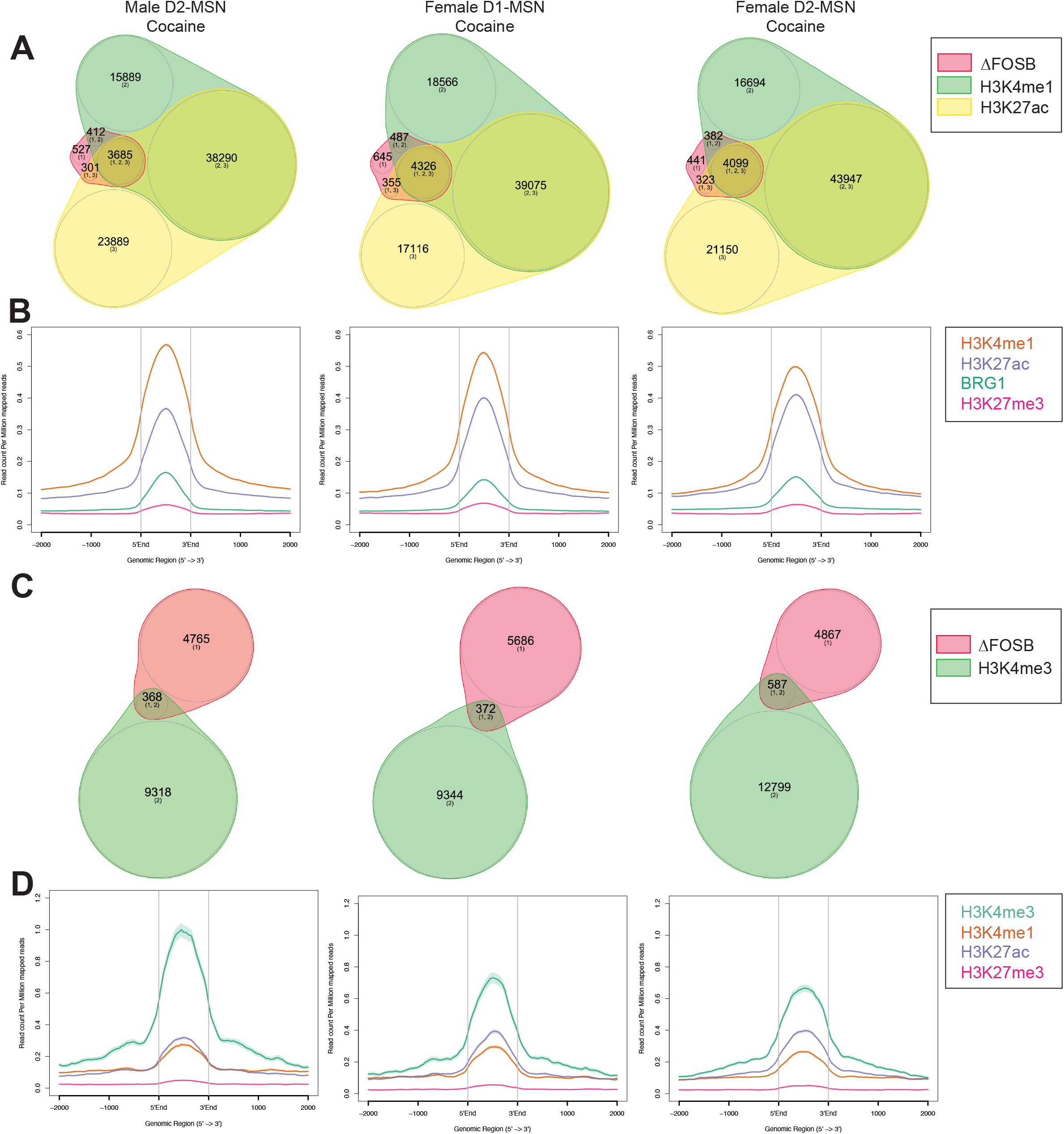
Overlap of ΔFOSB binding sites with BRG1 and histone marks in male D2 MSNs and female D1 and D2 MSNs. (A) Venn diagram analysis of called H3K4me1, H3K27ac, and ΔFOSB peaks. (B) Coverage of BRG1 and histone marks at called ΔFOSB+ / H3K4me1+ / H3K27ac+ peaks. (C) Venn diagram analysis of called H3K4me3 and ΔFOSB peaks. (D) Coverage of histone marks at called ΔFOSB peaks.

**Figure S4.**
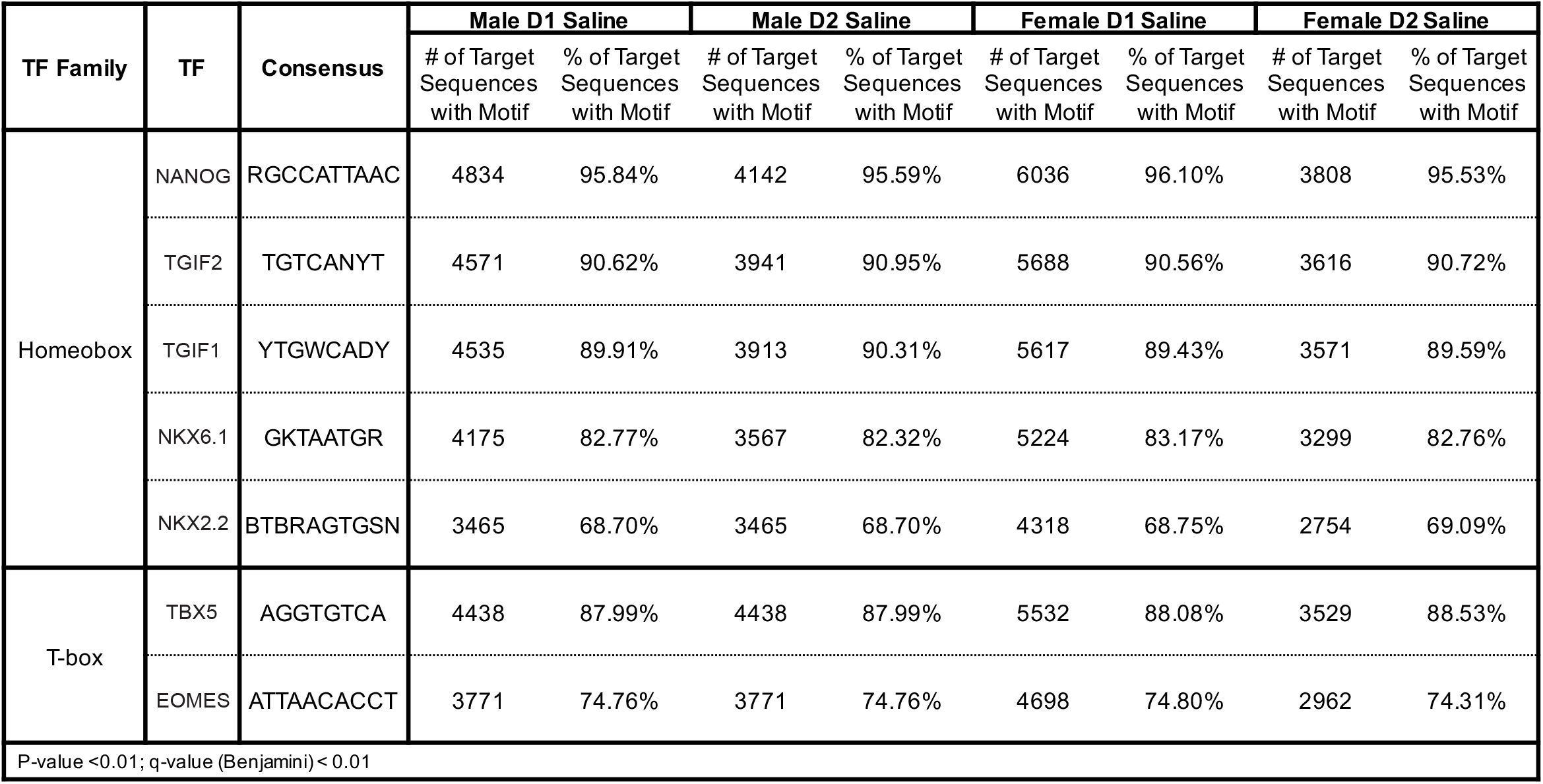
*In silico* motif analyses of loci with ΔFOSB-bound AP1 sites in NAc of male and female saline groups.

## References

1 Nestler, E. J. Review. Transcriptional mechanisms of addiction: role of DeltaFosB. Philos Trans R Soc Lond B Biol Sci 363, 3245–3255, doi:10.1098/rstb.2008.0067 (2008).

2 Robison, A. J. & Nestler, E. J. DeltaFOSB: A Potentially Druggable Master Orchestrator of Activity-Dependent Gene Expression. ACS Chem Neurosci 13, 296–307, doi:10.1021/acschemneuro.1c00723 (2022).

3 Hiroi, N. et al. FosB mutant mice: loss of chronic cocaine induction of Fos-related proteins and heightened sensitivity to cocaine’s psychomotor and rewarding effects. Proc Natl Acad Sci U S A 94, 10397–10402, doi:10.1073/pnas.94.19.10397 (1997).

4 Lobo, M. K. & Nestler, E. J. The striatal balancing act in drug addiction: distinct roles of direct and indirect pathway medium spiny neurons. Front Neuroanat 5, 41, doi:10.3389/fnana.2011.00041 (2011).

5 Calipari, E. S. et al. In vivo imaging identifies temporal signature of D1 and D2 medium spiny neurons in cocaine reward. Proc Natl Acad Sci U S A 113, 2726–2731 (2016), doi: 10.1073/pnas.1521238113.

6 Lobo, M. K. et al. DeltaFosB induction in striatal medium spiny neuron subtypes in response to chronic pharmacological, emotional, and optogenetic stimuli. J Neurosci 33, 18381–18395, doi:10.1523/JNEUROSCI.1875-13.2013 (2013).

7 Grueter, B. A., Robison, A. J., Neve, R. L., Nestler, E. J. & Malenka, R. C. FosB differentially modulates nucleus accumbens direct and indirect pathway function. Proc Natl Acad Sci U S A 110, 1923–1928, doi:10.1073/pnas.1221742110 (2013).

8 McClung, C. A. & Nestler, E. J. Regulation of gene expression and cocaine reward by CREB and DeltaFosB. Nat Neurosci 6, 1208–1215, doi:10.1038/nn1143 (2003).

9 Lardner, C. K. et al. Gene-Targeted, CREB-Mediated Induction of DeltaFosB Controls Distinct Downstream Transcriptional Patterns Within D1 and D2 Medium Spiny Neurons. Biol Psychiatry 90, 540–549, doi:10.1016/j.biopsych.2021.06.017 (2021).

10 Renthal, W. et al. Genome-wide analysis of chromatin regulation by cocaine reveals a role for sirtuins. Neuron 62, 335–348, doi:10.1016/j.neuron.2009.03.026 (2009).

11 Skene, P. J., Henikoff, J. G. & Henikoff, S. Targeted in situ genome-wide profiling with high efficiency for low cell numbers. Nat Protoc 13, 1006–1019, doi:10.1038/nprot.2018.015 (2018).

12 Mechta-Grigoriou, F., Gerald, D. & Yaniv, M. The mammalian Jun proteins: redundancy and specificity. Oncogene 20, 2378–2389, doi:10.1038/sj.onc.1204381 (2001).

13 Perrotti, L. I. et al. Distinct patterns of DeltaFosB induction in brain by drugs of abuse. Synapse 62, 358–369, doi:10.1002/syn.20500 (2008).

14 Winstanley, C. A. et al. DeltaFosB induction in orbitofrontal cortex mediates tolerance to cocaine-induced cognitive dysfunction. J Neurosci 27, 10497–10507 (2007), doi: 10.1523/JNEUROSCI.2566-07.2007.

15 Ohnishi, Y. N. et al. Generation and validation of a <em>floxed FosB</em> mouse line. bioRxiv, 179309, doi:10.1101/179309 (2017).

16 Walker, D. M. et al. Sex-Specific Transcriptional Changes in Response to Adolescent Social Stress in the Brain’s Reward Circuitry. Biol Psychiatry 91, 118–128, doi:10.1016/j.biopsych.2021.02.964 (2022).

17 Kyrchanova, O. & Georgiev, P. Mechanisms of Enhancer-Promoter Interactions in Higher Eukaryotes. Int J Mol Sci 22, doi:10.3390/ijms22020671 (2021).

18 Heintzman, N.D. et al. Histone modifications at human enhancers reflect global cell-type-specific gene expression. Nature 459, 108–112 (2009), doi.org/10.1038/nature07829

19 Herz, H.-M. et al. Enhancer-associated H3K4 monomethylation by Trithorax-related, the Drosophila homolog of mammalian Mll3/Mll4. Genes Dev. 26, 2604–2620 (2012), doi: 10.1101/gad.201327.112

20 Zabidi MA, et al. Enhancer-core-promoter specificity separates developmental and housekeeping gene regulation. Nature 518, 556–559 (2015), doi: 10.1038/nature13994.

21 Calo, E. & Wysocka, J. Modification of enhancer chromatin: what, how, and why? Mol Cell 49, 825–837, doi:10.1016/j.molcel.2013.01.038 (2013).

22 Mathur, R. & Roberts, C. W. M. SWI/SNF (BAF) Complexes: Guardians of the Epigenome. Annual Review of Cancer Biology 2, 413–427, doi:10.1146/annurev-cancerbio-030617-050151 (2018).

23 Ohnishi, Y. H. et al. PSMC5, a 19S Proteasomal ATPase, Regulates Cocaine Action in the Nucleus Accumbens. PLoS One 10, e0126710, doi:10.1371/journal.pone.0126710 (2015).

24 Kumar, A. et al. Chromatin remodeling is a key mechanism underlying cocaine-induced plasticity in striatum. Neuron 48, 303–314, doi:10.1016/j.neuron.2005.09.023 (2005).

25 Zachariou, V. et al. An essential role for DeltaFosB in the nucleus accumbens in morphine action. Nat Neurosci 9, 205–211, doi:10.1038/nn1636 (2006).

26 Renthal, W. et al. Delta FosB mediates epigenetic desensitization of the c-fos gene after chronic amphetamine exposure. J Neurosci 28, 7344–7349, doi:10.1523/JNEUROSCI.1043-08.2008 (2008).

27 McClung, C. A. et al. DeltaFosB: a molecular switch for long-term adaptation in the brain. Brain Res Mol Brain Res 132, 146–154, doi:10.1016/j.molbrainres.2004.05.014 (2004).

28 Corbett, B. F. et al. DeltaFosB Regulates Gene Expression and Cognitive Dysfunction in a Mouse Model of Alzheimer’s Disease. Cell Rep 20, 344–355, doi:10.1016/j.celrep.2017.06.040 (2017).

29 Robison, A. J. et al. Fluoxetine epigenetically alters the CaMKIIalpha promoter in nucleus accumbens to regulate DeltaFosB binding and antidepressant effects. Neuropsychopharmacology 39, 1178–1186, doi:10.1038/npp.2013.319 (2014).

30 Ferguson, D. et al. Essential role of SIRT1 signaling in the nucleus accumbens in cocaine and morphine action. J Neurosci 33, 16088–16098, doi:10.1523/JNEUROSCI.1284-13.2013 (2013).

31 Chen, J. et al. Induction of cyclin-dependent kinase 5 in the hippocampus by chronic electroconvulsive seizures: role of [Delta]FosB. J Neurosci 20, 8965–8971 (2000).

32 Nestler, E. J., Barrot, M. & Self, D. W. DeltaFosB: a sustained molecular switch for addiction. Proc Natl Acad Sci U S A 98, 11042–11046, doi:10.1073/pnas.191352698 (2001).

33 Maze, I. et al. Essential role of the histone methyltransferase G9a in cocaine-induced plasticity. Science 327, 213–216, doi:10.1126/science.1179438 (2010).

34 Vierbuchen, T. et al. AP-1 Transcription Factors and the BAF Complex Mediate Signal-Dependent Enhancer Selection. Mol Cell 68, 1067–1082 e1012, doi:10.1016/j.molcel.2017.11.026 (2017).

35 Kornblihtt, A. R. et al. Alternative splicing: a pivotal step between eukaryotic transcription and translation. Nat Rev Mol Cell Biol 14, 153–165, doi:10.1038/nrm3525 (2013).

36 Naftelberg, S., Schor, I. E., Ast, G. & Kornblihtt, A. R. Regulation of Alternative Splicing Through Coupling with Transcription and Chromatin Structure. Annual Review of Biochemistry 84, 165–198, doi:10.1146/annurev-biochem-060614-034242 (2015).

37 Xu, S. J. et al. Chromatin-mediated alternative splicing regulates cocaine-reward behavior. Neuron 109, 2943–2966 e2948, doi:10.1016/j.neuron.2021.08.008 (2021).

38 Feng, J. et al. Chronic cocaine-regulated epigenomic changes in mouse nucleus accumbens. Genome Biol 15, R65, doi:10.1186/gb-2014-15-4-r65 (2014).

39 Ma, X., Ezer, D., Adryan, B. & Stevens, T. J. Canonical and single-cell Hi-C reveal distinct chromatin interaction sub-networks of mammalian transcription factors. Genome Biol 19, 174, doi:10.1186/s13059-018-1558-2 (2018).

40 Wei, C. L. et al. A global map of p53 transcription-factor binding sites in the human genome. Cell 124, 207–219 (2006), doi: 10.1016/j.cell.2005.10.043.

41 Jägle, S. et al. SNAIL1-mediated downregulation of FOXA proteins facilitates the inactivation of transcriptional enhancer elements at key epithelial genes in colorectal cancer cells. PLoS Genet 13, e1007109 (2017), doi: 10.1371/journal.pgen.1007109.

42 Wu, Y. et al. Profiling transcription factor activity dynamics using intronic reads in time-series transcriptome data. PLoS Comput Biol 18, e1009762 (2022), doi: 10.1371/journal.pcbi.1009762.

43 Smemo, S. et al. Obesity-associated variants within FTO form long-range functional connections with IRX3. Nature 507, 371–375 (2014).

44 Hiroi, N. et al. Essential role of the fosB gene in molecular, cellular, and behavioral actions of chronic electroconvulsive seizures. J Neurosci 18, 6952–6962 (1998).

45 Jorissen, H. J. et al. Dimerization and DNA-binding properties of the transcription factor DeltaFosB. Biochemistry 46, 8360–8372, doi:10.1021/bi700494v (2007).

46 Eagle, A. L. et al. Experience-dependent induction of hippocampal ΔFosB controls learning. J Neurosci 35, 13773–13783 (2015), doi: 10.1523/JNEUROSCI.2083-15.2015.

47 You, J. C. et al. Epigenetic suppression of hippocampal calbindin-D28k by DeltaFosB drives seizure-related cognitive deficits. Nat Med 23, 1377–1383, doi:10.1038/nm.4413 (2017).

48 Manning, C. E. et al. Hippocampal Subgranular Zone FosB Expression Is Critical for Neurogenesis and Learning. Neuroscience 406, 225–233, doi:10.1016/j.neuroscience.2019.03.022 (2019).

49 Eagle, A. L. et al. Circuit-specific hippocampal ΔFosB underlies resilience to stress-induced social avoidance. Nat Commun 11, 4484 (2020), doi: 10.1038/s41467-020-17825-x.

50 Wang, Y. et al. Small molecule screening identifies regulators of the transcription factor DeltaFosB. ACS Chem Neurosci 3, 546–556, doi:10.1021/cn3000235 (2012).

51 Chen, X., Cho, K., Singer, B. H. & Zhang, H. The nuclear transcription factor PKNOX2 is a candidate gene for substance dependence in European-origin women. PLoS One 6, e16002, doi:10.1371/journal.pone.0016002 (2011).

52 Chen, I. C. et al. CUX2, BRAP and ALDH2 are associated with metabolic traits in people with excessive alcohol consumption. Sci Rep 10, 18118, doi:10.1038/s41598-020-75199-y (2020).

53 Li, M. D. & Burmeister, M. New insights into the genetics of addiction. Nat Rev Genet 10, 225–231, doi:10.1038/nrg2536 (2009).

54 Martin, M. Cutadapt removes adapter sequences from high-throughput sequencing reads. 2011 17, 3, doi:10.14806/ej.17.1.200 (2011).

55 Langmead, B. & Salzberg, S. L. Fast gapped-read alignment with Bowtie 2. Nat Methods 9, 357–359, doi:10.1038/nmeth.1923 (2012).

56 Danecek, P. et al. Twelve years of SAMtools and BCFtools. Gigascience 10, doi:10.1093/gigascience/giab008 (2021).

57 Robinson, J. T., Thorvaldsdottir, H., Wenger, A. M., Zehir, A. & Mesirov, J. P. Variant Review with the Integrative Genomics Viewer. Cancer Res 77, e31–e34, doi:10.1158/0008-5472.CAN-17-0337 (2017).

58 Thorvaldsdottir, H., Robinson, J. T. & Mesirov, J. P. Integrative Genomics Viewer (IGV): high-performance genomics data visualization and exploration. Brief Bioinform 14, 178–192, doi:10.1093/bib/bbs017 (2013).

59 Robinson, J. T. et al. Integrative genomics viewer. Nat Biotechnol 29, 24–26, doi:10.1038/nbt.1754 (2011).

60 Ramirez, F. et al. deepTools2: a next generation web server for deep-sequencing data analysis. Nucleic Acids Res 44, W160–165, doi:10.1093/nar/gkw257 (2016).

61 Zhang, Y. et al. Model-based analysis of ChIP-Seq (MACS). Genome Biol 9, R137, doi:10.1186/gb-2008-9-9-r137 (2008).

62 Li, Q., Brown, J. B., Huang, H. & Bickel, P. J. Measuring reproducibility of high-throughput experiments. The Annals of Applied Statistics 5, 1752-1779, 1728 (2011).

63 Quinlan, A. R. & Hall, I. M. BEDTools: a flexible suite of utilities for comparing genomic features. Bioinformatics 26, 841–842, doi:10.1093/bioinformatics/btq033 (2010).

64 Yu, G., Wang, L. G. & He, Q. Y. ChIPseeker: an R/Bioconductor package for ChIP peak annotation, comparison and visualization. Bioinformatics 31, 2382–2383, doi:10.1093/bioinformatics/btv145 (2015).

65 Ross-Innes, C. S. et al. Differential oestrogen receptor binding is associated with clinical outcome in breast cancer. Nature 481, 389–393, doi:10.1038/nature10730 (2012).

66 Heinz, S. et al. Simple combinations of lineage-determining transcription factors prime cis-regulatory elements required for macrophage and B cell identities. Mol Cell 38, 576–589, doi:10.1016/j.molcel.2010.05.004 (2010).

67 Perez-Silva, J. G., Araujo-Voces, M. & Quesada, V. nVenn: generalized, quasi-proportional Venn and Euler diagrams. Bioinformatics 34, 2322–2324, doi:10.1093/bioinformatics/bty109 (2018).

68 Shen, L., Shao, N., Liu, X. & Nestler, E. ngs.plot: Quick mining and visualization of next-generation sequencing data by integrating genomic databases. BMC Genomics 15, 284, doi:10.1186/1471-2164-15-284 (2014).

69 Carpenter, M. D. et al. Nr4a1 suppresses cocaine-induced behavior via epigenetic regulation of homeostatic target genes. Nat Commun 11, 504, doi:10.1038/s41467-020-14331-y (2020).

